# Dietary restriction induces post-transcriptional regulation of longevity genes

**DOI:** 10.1101/563296

**Authors:** Jarod A. Rollins, Santina S. Snow, Pankaj Kapahi, Aric N. Rogers

## Abstract

Dietary restriction increases lifespan through adaptive changes in gene expression. To understand more about these changes, we analyzed the transcriptome and translatome of *C. elegans* subjected to dietary restriction. Transcription of muscle regulatory and structural genes increased, while increased expression of amino acid metabolism and neuropeptide signaling genes was controlled at the level of translation. Evaluation of post-transcriptional regulation identified putative roles for RNA binding proteins, RNA editing, microRNA, alternative splicing, and nonsense mediated decay in response to nutrient limitation. Using RNA interference, we discovered several differentially expressed genes that regulate lifespan. We also found a compensatory role for translational regulation, which offsets dampened expression of a large subset of transcriptionally downregulated genes. Furthermore, 3’ UTR editing and intron retention increase under dietary restriction and correlate with diminished translation, while trans-spliced genes are refractory to reduced translation efficiency compared to messages with the native 5’ UTR. Finally, we find that *smg-6* and *smg-7*, which are genes governing selection and turnover of nonsense mediated decay targets, are required for increased lifespan under dietary restriction.

## Introduction

Methods of dietary restriction (DR) that increase lifespan involve restricting nutrients without causing malnutrition. Many forms of DR exist, including restriction of calories, specific macronutrients or micronutrients, and the timing of access to food (Greer and Brunet 2009; Honjoh et al. 2009, 1). Along with extended longevity, DR increases resistance to stress and improves other parameters of health from yeast to mammals (Mair and Dillin 2008), indicating that physiological responses to DR are evolutionary conserved. It is not surprising, then, that much attention has focused on understanding how animals adapt to DR. Identifying genetic programs that control the health benefits of DR guides efforts to develop drug mimetics, which would replace extreme dietary interventions to increase healthy longevity. In order to support such efforts, investigators need to understand the genes involved in adapting to DR so they can determine which ones impart benefits associated with this regimen.

Several molecular pathways and cellular processes are important for the effects of DR, especially those involved in nutrient sensing and energy status (Kapahi et al. 2010; Guarente 2011; Kenyon 2005). For example, energy sensing via AMP kinase and a greater role for energy production through aerobic respiration, along with changes to DNA and chromatin, are among the adaptive changes in the response to DR (Apfeld, 2004, 2; Vellai *et al*, 2003). Highly conserved pathways shown to have roles in increased lifespan under DR include the insulin/insulin-like signaling (ILS) and mechanistic target of rapamycin (mTOR) pathways. In response to changes in nutrient availability, the ILS pathways modulate cellular processes and coordinate responses in different tissues through hormone signaling (Lin *et al*, 2001, 1), while the TOR pathway can modulate cellular responses by directly sensing nutrients within the cell (Rohde *et al*, 2001). Changes in transcription associated with these pathways has been pivotal in resolving connections to biological processes and identifying new targets involved in increased lifespan (Weindruch *et al*, 2001; Han & Hickey, 2005; Zeier *et al*, 2011; Palgunow *et al*, 2012). However, studies have shown that these pathways also direct translation (Long *et al*, 2002; Kimball *et al*, 1994), including selective translation of specific mRNA species (Thoreen *et al*, 2012; Rogers *et al*, 2011; Zid *et al*, 2009; McColl *et al*, 2010).

The need to account for different levels of gene regulation is exemplified in a number of high profile studies showing that mRNA fluctuations, in general, account for less than half of the variability in corresponding proteins from yeast to humans (Ghaemmaghami *et al*, 2003; de Godoy *et al*, 2008; Schwanhäusser *et al*, 2011; Wilhelm *et al*, 2014). Thus, a comprehensive understanding of gene expression remodeling that takes place under DR requires an approach that accounts for transcript abundance as well as the efficiency with which individual transcripts are used to synthesize new protein. This approach provides the basis for development of hypotheses and tests of mechanisms that control gene expression at different levels. For example, it allows for the identification of putative sequence motifs within mRNA (i.e., *cis*-regulatory elements) and corresponding miRNAs and RNA binding proteins (i.e., *trans-* acting factors) targeting these motifs to ascertain their roles in translational control.

Today, methods involving polysome profiling (Zong *et al*, 1999) and ribosome footprinting (Ingolia *et al*, 2009) provide information that can be used to help distinguish forms of post-transcriptional regulation. By isolating mRNA that is associated with actively translating ribosomes, polysome profiling allows the translation state of individual transcripts to be quantified between treatments. In comparison, ribsome footprinting, allows for the distribution of the ribosome along mRNA to be quantified by sequencing only the regions of transcripts protected from nuclease digestion by bound ribosomes. These technologies have been applied in yeast to characterize the role of 5’ untranslated regions (UTRs) in transcript stability and translatability (Ringnér & Krogh, 2005), to pinpoint exonic polymorphisms in mRNA that affect translation, and to analyze stress-induced changes in translation of specific transcripts (Lackner *et al*, 2012). Polysome profiling and ribosome footprinting have also been used to consider isoform specific translational regulation (Sterne-Weiler *et al*, 2013) and to assess selective mRNA translation during mTOR inhibition in cell culture (Olshen *et al*, 2013). These studies and others like them have started to resolve the importance of post-transcriptional mechanisms in regulating gene expression. However, there is still a paucity of information on the relationships that exist between genes controlled transcriptionally, translationally, or both in mediating the adaptive response to DR. Understanding these relationships is key to understanding how DR promotes healthy aging.

In the following study, *C. elegans* adult nematodes were subjected to a form of DR involving food dilution that results in robust lifespan increase (Chen *et al*, 2009). Subsequently, animals were assessed for gene expression changes according to total and polysome-associated mRNA to differentiate transcriptional and post-transcriptional inputs. mRNAs differentially associated with polysomes under DR were investigated *in silico* for characteristics that could explain changes in gene expression at this level. Genes transcriptionally and/or translationally downregulated under DR were screened using RNA interference (RNAi) for effects on lifespan under fully fed conditions. Results reveal novel genes and connections with forms of post-transcriptional regulation important for increased lifespan under DR.

## Results

### Transcriptional responses to DR implicate muscle and metabolic adaptation

Using food dilution (Chen *et al*, 2009) to impose DR starting at day one of adulthood extended median lifespan of *C. elegans* by approximately 40% (Figure 1A). To better understand the different inputs to gene expression, we analyzed total and translated (i.e., polysome-associated) mRNA after 4 days under DR or fully fed (<u>a</u>d libitum, or AL) conditions in adult nematodes. Using four biological replicates, total and translated mRNA from AL and DR worms were analyzed by next-generation sequencing (NGS; Figure 1B). In brief, 100 bp paired end reads were aligned to the *C. elegans* genome and reads were counted using HTseq (Figure 1C). EdgeR was used to normalize the dataset and filter out lowly expressed mRNA that could not be reliably quantified (See Materials and Methods), leaving 8,301 annotated coding genes in the analysis of differential expression. A heatmap representing expression across this dataset demonstrated that the biological replicates clustered together, that the expression dataset was normally distributed, and that several instances of differential expression existed between different groups (Figure EV1A).

**Figure 1.**
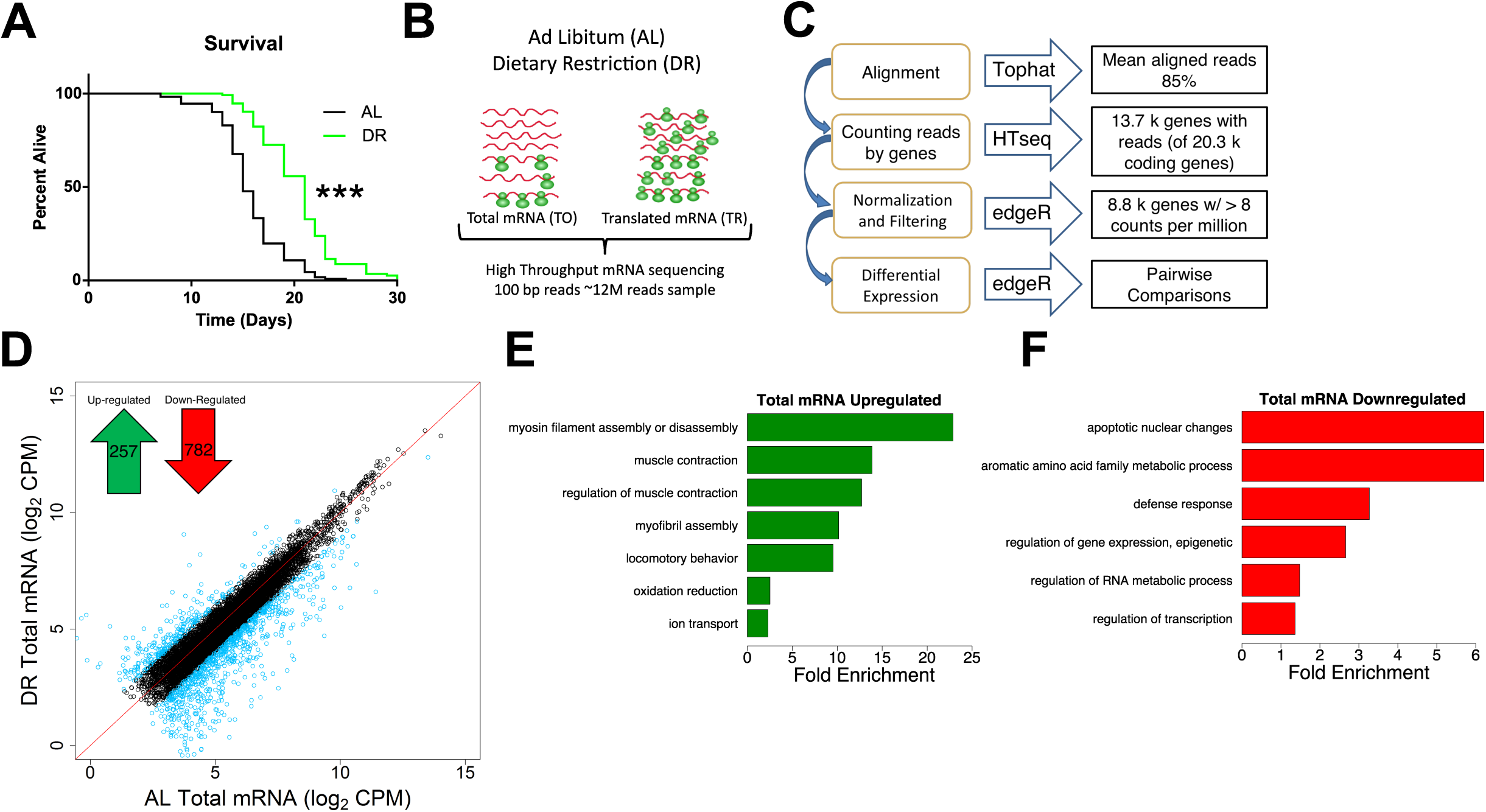
Transcriptional mRNA regulation under DR in *C. elegans*. **A)** Dietary restriction increased median lifespan by 40% compared to well fed conditions. p < 0.0001, n = 120, log-rank test. **B)** Schematic of experimental setup combining mRNA-seq and polysome profiling to quantify gene expression. At day one of adulthood, nematodes were placed on Ad libitum (AL) or DR plates for four days. Total (TO) and translated (TR) mRNA were extracted from whole worm populations in biological quadruplicate and subjected to high throughput sequencing. **C)** Schematic of deep sequencing analysis. Reads were aligned to the *C. elegans* reference genome WS220 using Tophat and were counted using HTseq. Normalization, filtration, and differential expression were performed by edgeR. A total of 8,301 genes were quantified. **D)** Protein coding genes are differentially expressed under DR on the transcriptional level as determined using reads from total mRNA. Each circle represents expression for a specific gene. Black circles plotted near the red line represent genes with the same or similar expression between diets, while blue circles represent genes with two-fold or greater change. Units are log_2_ values of counts aligned per million reads sequenced (CPM). Inset, 257 protein coding genes were up-regulated and 782 were down-regulated significantly (FDR < 0.05) 2-fold or more under DR. **E)** and **F)** show biological GO terms significantly (* P < 0.05, modified Fisher’s Exact) enriched among genes transcriptionally upregulated or downregulated under DR, respectively.

To compare total mRNA expression between AL and DR conditions, we identified genes that were significantly changed two-fold or more between diets (FDR < 0.05; Figure 1D). Of these, 257 were upregulated under DR and 782 were downregulated (Table EV1). Gene ontological (GO) analysis was then used to determine whether particular biological processes were represented among differentially expressed genes. Upregulated genes were enriched for processes related to myosin and myofibril assembly, muscle contraction, locomotion, oxidation/reduction and ion transport (Figure 1E and Table EV2), whereas downregulated genes were enriched for biological processes related to apoptosis, amino acid metabolism, defense response, epigenetic regulation, RNA metabolism, and transcription (Figure 1F and Table EV2).

Among the upregulated genes, expression of several are known to be critical for age-related health and longevity. Like age-related sarcopenia in humans, locomotion and the ability to transduce force generated by muscles is lost with age in *C. elegans* (Rollins *et al*, 2017; Hahm *et al*, 2015). Under DR, the genes encoding thick filaments (*unc-54* and *myo-1*) and movement-coordinating troponins (*pat-10*, *tni-3* and *tnt-4*) of muscle sarcomeres were upregulated (Table EV2), suggesting that they may be involved in the maintenance of locomotion with age in DR restricted animals. In line with this, *C. elegans* mutants with defective *unc-54* activity have reduced capacity to generate movement force (Várkuti *et al*, 2012). Additionally, overexpression of *pat-10* leads to increased actin filament stability, resistance to protein unfolding stress and lifespan (Baird *et al*, 2014). Among the genes with roles in oxidation reduction were members of the cytochrome P450 family *(cyp-34A9, cyp-35A2, cyp-33C3,* and *cyp-34A4*). Cytochrome P450 isozymes can serve in the metabolism of xenobiotics and other toxins (Ding & Kaminsky, 2003; Lindblom & Dodd, 2006) which might otherwise impinge on longevity.

Defense response genes were downregulated, including those encoding neuropeptide antimicrobials (e.g., *npl-29* and *npl-31*) in addition to *tol-1*, the only Toll-like receptor in *C. elegans*. Although efficacy of defense responses is usually improved in DR animals, diminished expression of these genes may be due to reduced microbe sensing resulting from the form of DR itself, which involves dilution of their bacterial food source. The transcription factor GO term included regulators of embryonic development, which were expected to be decreased given the negative effect of DR on reproduction. These included nuclear hormone receptors (e.g., *nhr-2* and *nhr-112*), cyclin-dependent kinase 8 (*cdk-8*), a homeobox factor (*ceh-40)*, and a HAND bHLH factor important for tissue differentiation (*hnd-1)*, among others (Table EV2). The heat shock transcription factor gene *hsf-1* was also downregulated, presumably because of globally reduced protein synthesis, which requires *hsf-1*-dependent chaperones to direct the folding of nascent proteins.

In a previous study, transcript levels were analyzed in *C. elegans* subjected to DR based on intermittent fasting (Honjoh *et al*, 2009). We compared our list of differentially regulated genes from bacterial dilution to those obtained by Honjoh et al. (2009) using gene set enrichment analysis (GSEA, Figure EV1B). This analysis demonstrated a significant (FDR < 0.001) enrichment of both up-and downregulated genes between the two datasets despite differences in dietary regimes and quantification methods (i.e. NGS vs microarray). To gain insight into the transcriptional differences between DR by food dilution and intermittent fasting, differentially expressed genes in common between DR methods and those unique to food dilution were subjected to GO analysis. Terms related to oxidation/reduction and muscle contraction were upregulated in common among both forms of DR (Table EV2). Among downregulated genes, terms in common pertained to defense and immune response. However, the term “regulation of transcription” was unique to downregulated genes in the food dilution DR dataset (Table EV2). Together, results demonstrate that many of the same processes regulated transcriptionally by intermittent fasting are also regulated by food dilution in *C. elegans*.

In order to gain a better understanding of which responses to DR are species specific and which are conserved among different species, we compared our differential expression results with a meta-analysis performed by Han and Hickey (2005). In the meta-analysis, they compared differently expressed genes in mouse, rat, pig, monkey, yeast, and fly subjected to DR. Interestingly, their study showed a lack of overlap in expression changes of specific genes in common to all these species. However, GO analysis did point to the conservation of biological processes involved in adaptive responses to DR like those observed in the present study. Specifically, the meta-analysis reported the enrichment of genes annotated with the term metabolism among all animals listed. In addition, there was a high level of overlap for terms related to stress/immune response and regulation of transcription that were the same or similar in *C. elegans* (Figure 1F). These similarities imply evolutionary conservation of transcriptional responses to DR at the level of biological processes.

### Translation frequently offsets transcriptional downregulation under DR

To differentiate transcriptional and translational regulation of gene expression, we analyzed the relative abundance of the 8,301 quantified genes within the total and polysome-associated mRNA pools. Under AL conditions, 82% of genes had similar abundance between pools. The remaining 18% showed a 2-fold or greater difference between pools and were considered to be translationally regulated (Figure 2A). Thus, genes with greater relative abundance among polysomes than total mRNA were considered to be translationally promoted, while those with reduced polysomal association were considered translationally suppressed. In contrast to AL conditions, 34% of genes were translationally regulated under DR (Figure 2B). This is reflected by reduced correlation between total and translated mRNA pools (r = 0.88 for AL versus 0.83 for DR, p < 0.0001, t-test of Fisher r-z transformed coefficients). Additionally, the distribution of changes in gene expression between the total and translated fractions was significantly different under DR compared to AL (p < 2.2e-16, two-sample Kolmogorov-Smirnov test, Figure 2C). As seen in the Venn diagram inset in Figure 2C, the majority of selectively translated genes under AL were also subject to translational regulation under DR. Thus, a common set of genes are translationally regulated under both conditions, while an additional set of genes becomes translationally regulated under DR.

**Figure 2.**
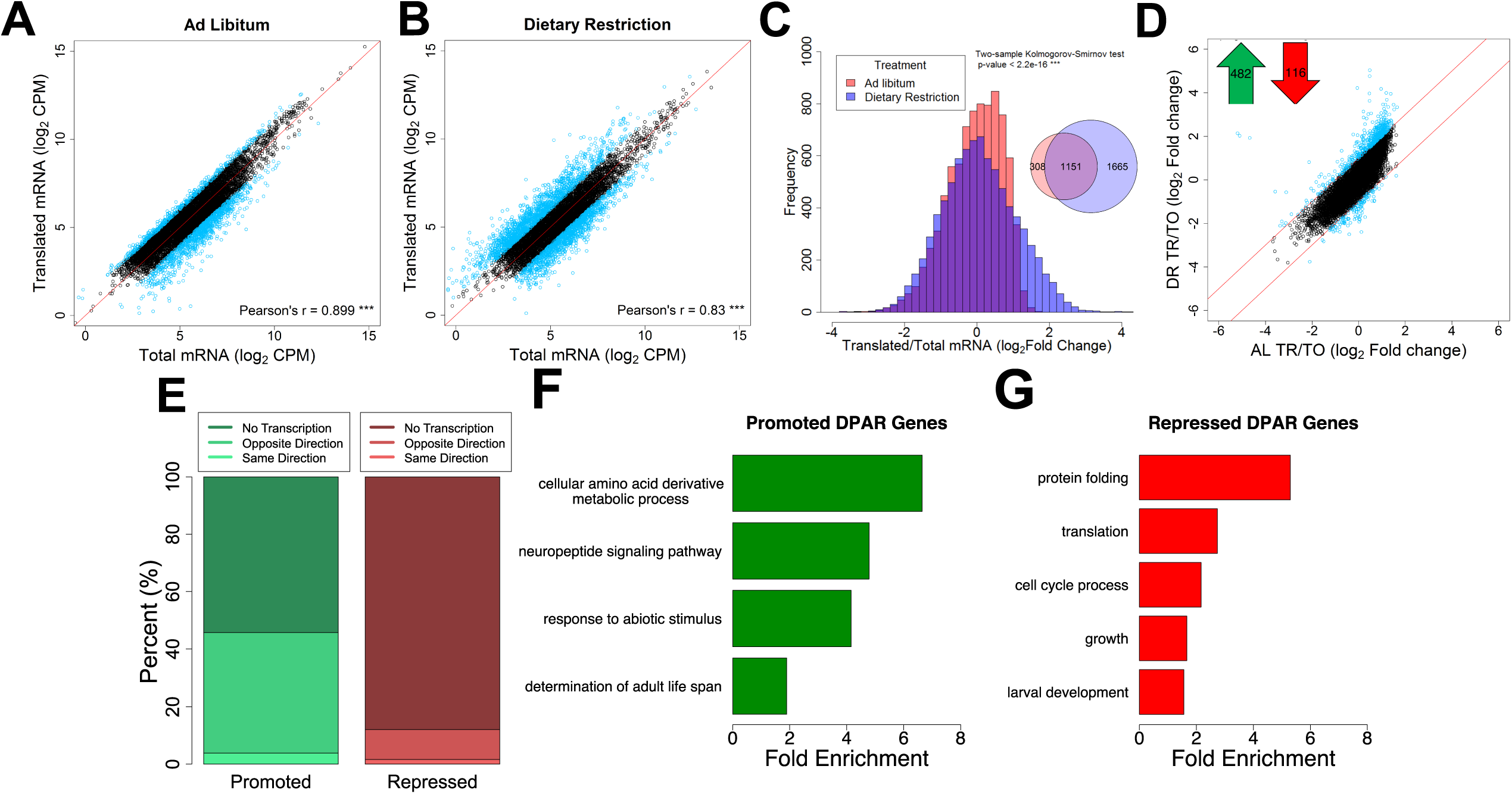
DR modulates selective translation in *C. elegans*. Gene abundance between fractions was well correlated under **A)** ad libitum and **B)** dietary restriction (p-val < 0.001, Pearson’s correlation of 0.899 and 0.830, respectively). Black circles represent genes that showed no significant difference (FDR > 0.05) between total and translated mRNA. Blue circles represent genes with two-fold or greater change. A reference line with a slope of 1 is shown in red. Units are counts aligned per million sequenced (CPM). **C)** Distribution of gene abundance fold changes between translated and total mRNA for AL (red) and DR (blue) worms. The distribution of changes was significantly different between diets (Two-sample Kolmogorov-Smirnov test, p < 2.2e-16). **D)** Translational promotion and repression of genes between diets were calculated using the DPAR. Black circles represent genes with similar translation:transcription expression ratios between diets. Blue circles represent genes with two-fold or greater DPAR. Red lines delineate the two-fold change cutoff. Inset, 482 genes were translationally promoted and 116 were translationally repressed. **E)** Percent of translationally promoted or repressed DPAR genes in which changes in the total fraction were either in the same direction (light hue), opposite direction (medium hue), or were not significant (dark hue; Log2 FC >2, FDR < 0.05) Biological GO terms associated with translationally **F)** Promoted or **G)** Repressed DPAR genes under DR. P < 0.05, modified Fisher’s Exact test. Functionally similar GO terms from identical groups of genes were manually removed.

To characterize the predominant modes of gene regulation that occur under DR without bias, we clustered genes that only become translationally regulated under DR (1,665 in all, see Venn diagram in Figure 2C) into groups with similar expression profiles. Partitioning around medoids was used to cluster genes into the optimal number of groups using the gap statistic (Tibshirani, Walther, and Hastie 2001). These genes were optimally clustered into three groups as seen in a PCA plot that explained 97.6% of the dataset variance (Figure EV2A). The relative expressions of all 1,665 genes for total and translated mRNA under AL and DR are shown for each of the three clusters in Figure EV2B-D. The first cluster consisted of 530 genes exhibiting downregulated transcript levels under DR that were partly offset by positive selective translation (Figure EV2B). The second group contained 492 genes lacking transcriptional changes with modest negative translation changes under AL that became more negative under DR (Figure EV2C). The third cluster contained 643 genes that were essentially unchanged transcriptionally under both conditions but that become translationally promoted under DR (Figure EV2D). While other modes of regulation exist under DR, these clusters account for the vast majority of expression changes. Thus, it appears that there are three fundamental roles of translational regulation that occur under this DR regimen: to oppose transcriptional down regulation (Cluster 1), to reinforce translational suppression tendency observed under AL (Cluster 2), and to increase expression during DR in the absence of transcriptional changes (Cluster 3).

The analysis in Figure EV2 helps resolve patterns of expression for genes that only become translationally regulated subsequent to DR. In order to determine which genes exhibit net translation changes between conditions, we devised the Differential Polysome Association Ratio (DPAR). For a given gene, DPAR is calculated by taking the ratio of translated to total mRNA under DR and dividing by the same ratio under AL. A significant (FDR < 0.5) 2-fold or greater change in translation was observed for 482 translationally promoted genes and 116 translationally suppressed genes (Figure 2D and Table EV3). A threshold of twofold change in polysome association was used to focus the analysis on genes experiencing large changes in translational regulation and comes with the tradeoff of potentially missing smaller but still biologically relevant regulation. Interestingly, the tendency for translationally changed genes to be upregulated under DR was opposite of the tendency for transcriptionally changed genes, which were much more frequently downregulated (compare Figure 1D and 2D). This led us to investigate how often translation offsets transcriptional regulation under DR. Of the genes that were translationally promoted under DR, 58% lacked significant transcript level changes and 40% had significantly decreased transcript level changes, leaving very few genes that were both transcriptionally and translationally promoted (Figure 2E). Given that only 12.5% of quantifiable genes in our dataset displayed significant transcriptional regulation, the fact that 33% of translationally promoted genes were significantly downregulated at the transcript level was, itself, significant (p < 2.2e-16, Fisher’s exact test). Conversely, 90% of translationally downregulated genes showed no change in transcription. The frequent opposition between translation and transcription may help explain numerous studies showing disparate results between gene transcription and protein expression (Ghaemmaghami *et al*, 2003; Schwanhäusser *et al*, 2011; Wilhelm *et al*, 2014).

### Translational adaptation to DR involves downregulating anabolic processes and altering amino acid metabolism and neuropeptide signaling

We performed GO enrichment analysis of DPAR genes to determine which biological processes are candidates for DR-relevant adaptation through translation. Translationally promoted DPAR genes were enriched (p > 0.05) for processes related to amino acid metabolism, neuropeptide signaling, abiotic responses, and determination of adult lifespan (Figure 2F and Table EV4). One example is *gst-10*, a glutathione S-transferase for which overexpression has been shown to increase median longevity in *C. elegans* by 22% (Ayyadevara *et al*, 2005). Interestingly, translationally promoted DPAR genes *dod-19*, *dod-23*, and *mtl-2* were previously shown to be downregulated in *daf-2* ILS mutants and could further increase lifespan when knocked down in the *daf-2(mu150)* background (Murphy *et al*, 2003). In the case of *dod-19* and *dod-23*, their transcriptional down regulation under DR (>24-fold and >5-fold reduction, respectively) was far greater than their translational promotion (8-fold and 2-fold increase, respectively) which is likely to result in an overall reduction in new protein synthesis of these genes under DR, consistent with their negative roles in longevity. The *mtl-2* mutant has a lifespan similar to wild-type worms (Hughes & Stürzenbaum, 2007), which is in line with a context dependent role in longevity. Among the translationally suppressed genes there was an enrichment of biological terms related to protein folding, translation, cell cycle, growth and development (Figure 2G and Table EV4). These results are in agreement with effects of DR on limiting growth and decreasing global translation.

### RNAi screen of translationally regulated genes revealed novel DR longevity genes

As DR prolongs longevity, we tested whether inhibiting genes that are downregulated under food dilution increases lifespan in *C. elegans*. To avoid deleterious effects on growth, RNAi was initiated after development was complete. We included the top 100 transcriptionally downregulated genes and the top 95 translationally downregulated genes under DR using the Ahringer (Kamath *et al*, 2003; Fraser *et al*, 2000) and ORFeome (Rual *et al*, 2004) RNAi libraries. These genes are listed in Table EV5.

Three of the 100 transcriptionally suppressed genes resulted in a significant extension in median lifespan when targeted with RNAi: *fbxb-24, sdz-33,* and D1086.3 (Table 1). The genes *fbxb-24* and *sdz-33* encode proteins containing F-boxes, which have been implicated in mediating protein-protein interactions important for cell cycle regulation and signal transduction (Craig & Tyers, 1999). Little information is available for D1086.3, but it is the only gene both transcriptionally and translationally downregulated with a lifespan phenotype (Table EV5, Table 1). The list of screened genes also included *dod-19*, *dod-23*, and *clec-186*, which have been found in previous studies to increase lifespan when knocked down but which did not increase lifespan in the current study. *dod* genes are named for being <u>D</u>ownstream <u>o</u>f <u>D</u>AF-16, a forkhead box O transcription factor required for increased lifespan when the ILS pathway is attenuated (Murphy *et al*, 2003). These *dod* genes are downregulated when the ILS pathway is genetically restricted and targeting them with RNAi in *daf-2*(*mu150*) animals further increases the long lifespan associated with background (Murphy *et al*, 2003). Our results indicate that these *dod* genes may require muted ILS to effectively increase lifespan when they are targeted with RNAi. Knockdown of the innate immunity gene *clec-186* was shown to extend lifespan in *C. elegans* with increased sensitivity to RNAi (Hamilton *et al*, 2005), suggesting that the total or tissue-specific extent of suppression is a factor for its role in lifespan.

**Table 1.**
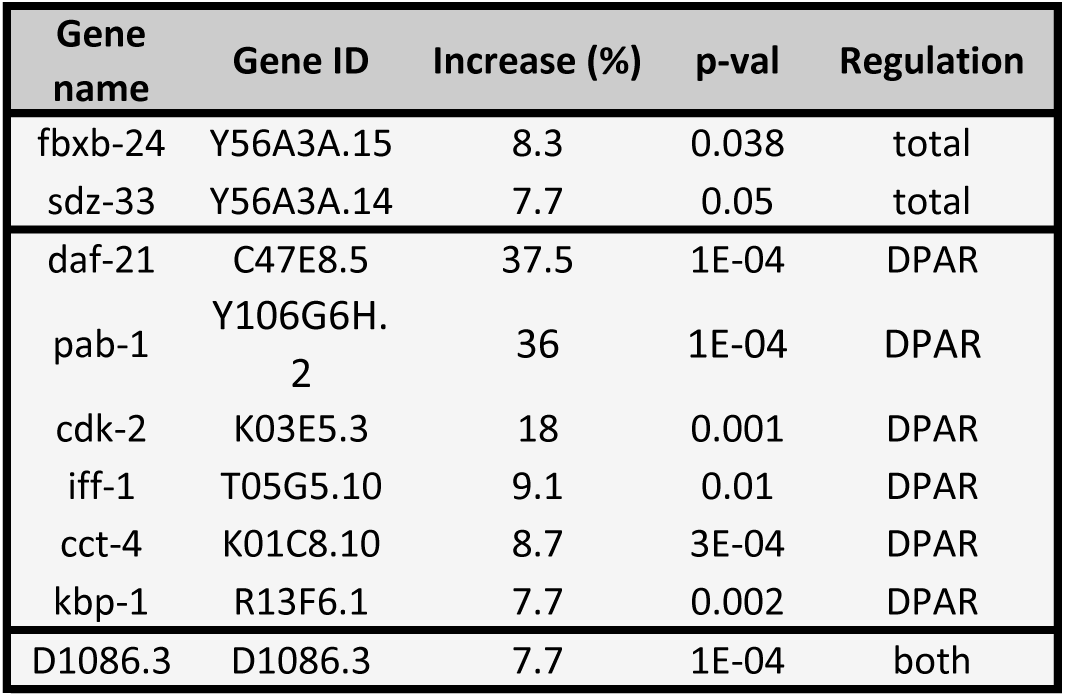
Genes resulting in lifespan extension when knock-downed in *C. elegans* using RNAi under well fed-conditions. The gene common name and gene ID are given as well as the percent increase in median lifespan as compared to control empty vector. The p-values are from a Log-rank (Mantel-Cox) test. Regulation is based on the gene expression in our RNA-seq dataset. DPAR – differential polysomal association ratio, meaning the gene was predominantly regulated post-transcriptionally under DR, total - the gene was predominantly regulated in the total fraction, meaning the gene was regulated transcriptionally, both – the gene evidenced regulation both transcriptionally and post-transcriptionally.

| Gene name | Gene ID | Increase (%) | p-val | Regulation |
| --- | --- | --- | --- | --- |
| fbxb-24 | Y56A3A.15 | 8.3 | 0.038 | total |
| sdz-33 | Y56A3A.14 | 7.7 | 0.05 | total |
| daf-21 | C47E8.5 | 37.5 | 1E-04 | DPAR |
| pab-1 | Y106G6H.2 | 36 | 1E-04 | DPAR |
| cdk-2 | K03E5.3 | 18 | 0.001 | DPAR |
| iff-1 | T05G5.10 | 9.1 | 0.01 | DPAR |
| cct-4 | K01C8.10 | 8.7 | 3E-04 | DPAR |
| kbp-1 | R13F6.1 | 7.7 | 0.002 | DPAR |
| D1086.3 | D1086.3 | 7.7 | 1E-04 | both |

Seven of the 95 downregulated DPAR genes increased lifespan when targeted for knockdown via RNAi, including D1086.3. The other six genes are *daf-21*, *cct-4*, *cdk-2*, *iff-1*, *pab-1,* and *kbp-1* (Table 1). The *daf-21* gene encodes the ortholog of the mammalian molecular chaperone HSP90. Despite its title as a heat shock protein, HSP90 is a pro-growth chaperone that inhibits activity of the heat shock transcription factor (Kijima *et al*, 2018). Lowering its expression was recently shown to increase lifespan in wild-type animals in a separate study investigating *C. elegans* orthologs of human genes that are differential expressed with age in blood (Sutphin *et al*, 2017). *cct-4* is a chaperonin required for SKN-1-dependent transcription of a putative glutathione-requiring prostaglandin D synthase, *gst-4* (Kahn *et al*, 2008). The gene *cdk-2* encodes for a cyclin-dependent kinase which has been recently implicated in regulating stress responses and longevity by inhibiting cell cycle machinery (Dottermusch *et al*, 2016). A germline specific translation initiation factor is encoded by *iff-1*, which was shown to regulate lifespan in the first genome wide functional screen for longevity genes (Hamilton *et al*, 2005). Another translation initiation factor, *pab-1* encodes a protein that binds the polyA tail of mRNA and is discussed more in the next section. The gene *kbp-1* is involved in the cell cycle by regulating kinetochore-microtubule binding (DeLuca & Salmon, 2004) and is necessary for larval development (Rual *et al*, 2004). To our knowledge, our findings for *kbp-1* and D1086.3 are the first to demonstrate that they play a role in regulating aging in *C. elegans*. Furthermore, these results demonstrate the importance of understanding the role of translational regulation of gene expression in longevity.

### Translationally suppressed genes under DR are frequently in operons

Analysis of the switch from AL to DR showed that translationally promoted genes were frequently associated with diminished transcript levels (Figure 2E and E2B), pointing to a possible link between transcription and translation in dietary adaptation. To address mechanisms governing translation, as well as a possible connection to transcriptional regulation, we investigated the roles of operons and *trans*-splicing. The genome of *C. elegans* contains operons consisting of two or more genes that are transcribed together but separated post-transcriptionally and translated individually (Zorio *et al*, 1994). We hypothesized that expression of genes of the same operon may require compensatory translational regulation in response to changes in nutrient availability. Using previously established operon annotations (Allen *et al*, 2011), we calculated the percent of genes in operons among all coding genes and compared this to the percent found among translationally promoted or suppressed DPAR genes (Figure 3A). Of the 482 promoted genes, a significantly smaller proportion (10%, p < 0.001, Fisher’s exact test) were organized in operons compared to 8,301 coding genes (18%) with quantifiable expression in our dataset. Conversely, the genes translationally suppressed contained a higher proportion of operonic genes (50%, p < 0.05, Fisher’s exact test). This observed enrichment suggested that the translational suppression observed under DR may serve to suppress the expression of one or more genes that are co-transcribed at levels higher than needed under DR. This pattern was inverted among the majority of translationally promoted genes, which tended not to be associated with operons (Figure 3A) and which, instead, tended to be associated with transcriptionally suppressed genes (Figure 2E). To gain a better understanding of this association, we addressed the known attributes of operonic and non-operonic genes that can influence their rate of translation.

**Figure 3.**
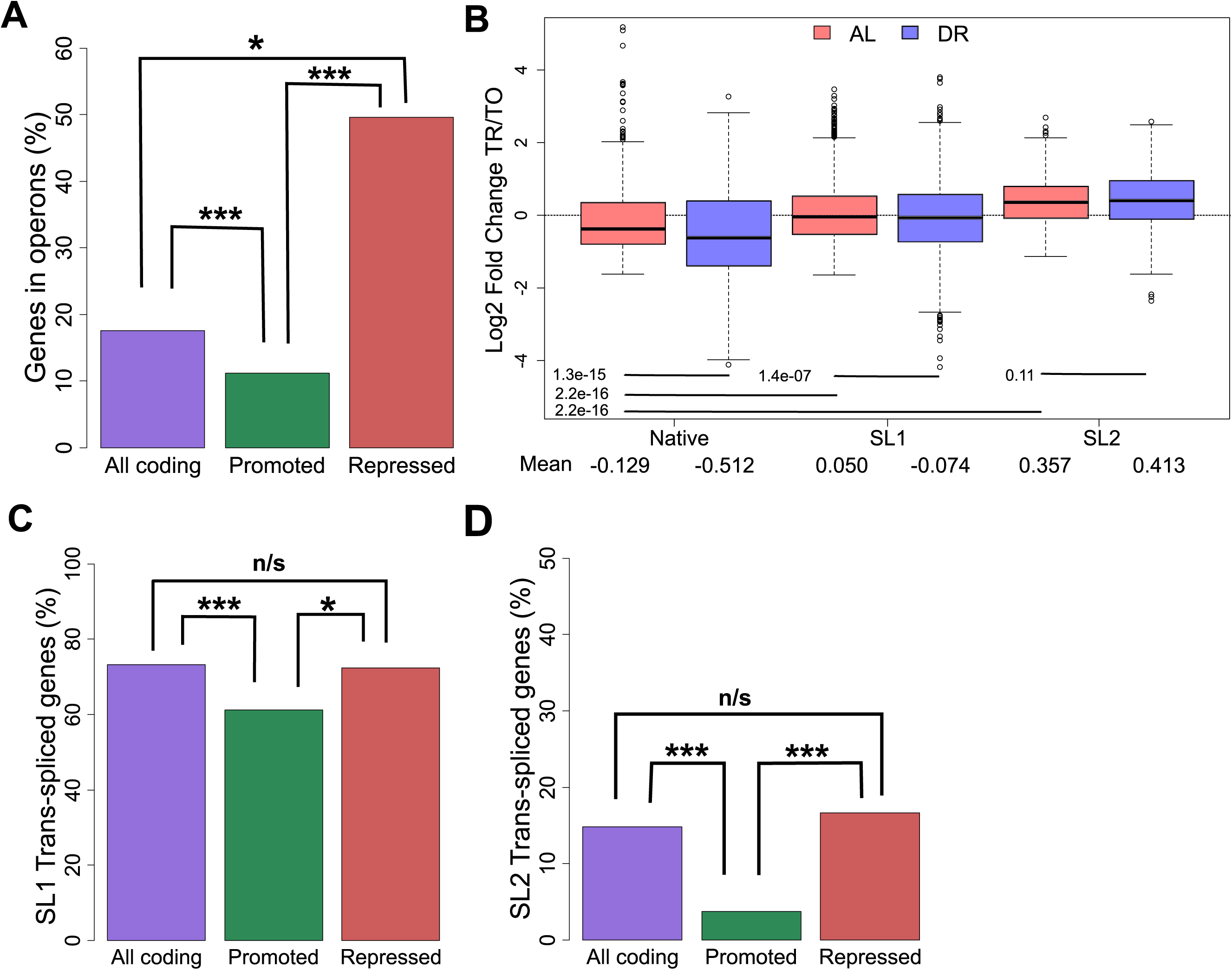
Operonic genes are enriched among genes that undergo translational repression in the switch to DR. **A)** Percent of operonic genes among all protein coding genes (purple), translationally promoted DPAR genes (green), or translationally repressed genes (red). **B)** Translational regulation of genes under AL (light red) or DR (blue). Genes are grouped by their predominant 5’UTR, which may be native or *trans*-spliced to SL1 or SL2. The mean Log2 fold change between translated and total fractions is given for AL and DR. Group means were compared using Wilcoxon rank sum test and the associated p-values are given. **C)** Percent of genes SL1 trans-spliced among all protein coding genes (purple), translationally promoted genes (green), or among translationally repressed genes (red). **D)** Percent of SL2 spliced genes among all protein coding genes, translationally promoted genes and translationally repressed genes under DR. TO – total fraction, TR – translated fraction. Asterisks denote significance (* p < 0.05, ** p < 0.01, *** p < 0.001) using Fisher’s exact test.

### *Trans-*spliced genes are refractory to diminished translation under DR compared to other genes

The 5’ UTR of many *C. elegans* mRNA are *trans-*spliced with one of a small number of short spliced leader (SL) sequences that replace much of the originally transcribed 5’ UTR. 5’ UTRs are important for regulating translational efficiency and *trans-*splicing has a positive effect on translation in unstressed *C. elegans* (Yang *et al*, 2017). Commonly, the first gene in an *C. elegans* operon is *trans-*spliced to SL1 and all downstream transcripts are *trans-*spliced to SL2 (Allen *et al*, 2011). For genes that are not in operons, 5’ UTRs are frequently *trans-*spliced to SL1, with around 30% of transcripts retaining the native 5’ UTR (Allen *et al*, 2011).

Using annotations of SL1 and SL2 spliced genes from Allen et al (2011), we calculated the average fold change in translational regulation of genes with *trans-*spliced or native 5’ UTRs. Under AL, our analysis confirmed that both SL1 and SL2 *trans-*spliced messages tend to be better translated than transcripts with native 5’ UTRs (Figure 3B), as previously reported (Yang *et al*, 2017). DR led to translational suppression of SL1 spliced genes compared to AL conditions (p=1.4e-7, Wilcoxon rank sum test with continuity correction). In contrast, the translation efficiency of SL2 spliced genes was unchanged under DR (p=0.11, Wilcoxon rank sum test with continuity correction). For both SL1 and SL2 genes, translation was preserved better in the transition from AL to DR compared to those annotated with native 5’ UTRs.

Approximately 73% of coding genes quantified in this study were annotated as SL1 *trans-*spliced (Figure 3C). In comparison, a significantly lower proportion of SL1 *trans-*spliced transcripts were among promoted DPAR genes (61%, p < 0.01, Fisher’s exact test) while a similar proportion was observed among repressed genes (63%, p > 0.05, Fisher’s exact test). A similar analysis for SL2 *trans-*spliced transcripts showed a significantly lower proportion among promoted DPAR genes (4%, Figure 3D, p < 0.001, Fisher’s exact test) but not among suppressed DPAR genes (17%, p > 0.05, Fisher’s exact test). Thus, whilst promoted DPAR genes were depleted among SL1 and SL2 transcripts, the depletion is modest and no change from the average distribution of trans-splicing was observed for repressed DPAR genes, suggesting that *trans-*splicing is unlikely to be a primary driver of differential translation among DPAR genes.

### DR increases 3’ UTR editing, which is associated with reduced translation

Within the RNA-seq dataset we found a number of putative adenosine RNA editing events with the potential to alter translational regulation. The modification of adenosine to inosine (A-to-I) is the most prevalent type of RNA editing in many species and is catalyzed by action of Adenosine Deaminase Acting On RNA (ADARs). Inosine is read as a guanosine (G) by cellular enzymes, which has the potential to change protein-coding regions and binding sites. The A-to-I modification may also change the secondary structure of non-coding regions. For some genes, like the glutamate receptor in mice, RNA editing is required to produce a functional protein (Wright & Vissel, 2012).

To determine the influence of DR on editing, we quantified A to I conversions occurring within each of four biological replicates for all test groups in our dataset. For an edit site to be included, we required a minimum editing frequency of 10%. Under AL, we detected an average of 125 putative editing sites in the total fraction, which was significantly decreased to an average of 70 in the translated fraction (Figure 4A), indicating that the occurrence of A-to-I editing is negatively associated with translation (p = 0.029, Wilcoxon rank sum test with continuity correction). DR amplified this difference, increasing putative editing sites to 154 with a concomitant decrease to 47 sites in the translated fraction (p = 0.021, Wilcoxon rank sum test with continuity correction). A similar pattern is observed with respect to the frequency of editing at these sites (Figure EV3A). The overlap of editing sites amongst the different experimental groups is shown in Figure 4B, which shows 39 sites common to all samples, but none that are exclusive to the translated mRNA under DR. Together, these results suggest that A-to-I editing may be used to effectively diminish expression of specific genes under DR.

**Figure 4.**
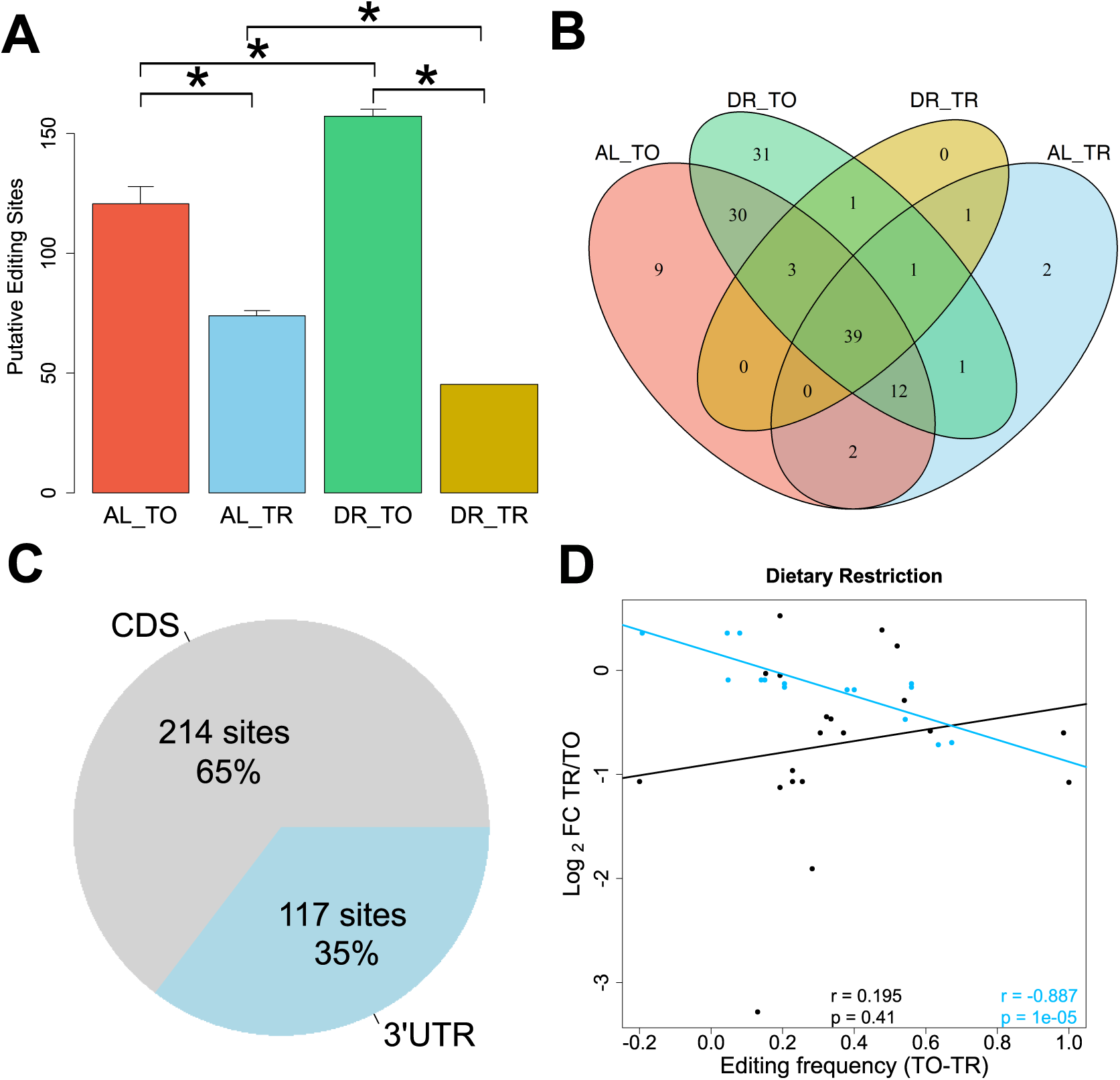
mRNA adenosine-to-inosine editing increases under DR. **A)** Average number of RNA edits in total and translated fractions across diets. RNA editing under DR is globally increased in the total fraction yet diminished in the translated fraction as compared to well fed controls. Error bars are s.e.m. * P < 0.05, Wilcoxon rank sum test, n=4. **B)** Overlap of editing sites present in at least 3 out of 4 all biological replicates in total and translated mRNA under AL or DR. **C)** Percentage of editing sites within the dataset located in 3’UTRs as annotated in UTRome and the coding sequence (CDS). **D)** Relationship between the change in editing frequency and polysome association under DR. Genes with edits occurring in their 3’ UTR are blue, edits occurring in the CDS are black. Regression line for 3’UTR edits is blue, line for CDS edits is black. Spearman correlation coefficient and associated p-value for 3’UTR edits and for CDS genes are given in blue and black, respectively. TO – total mRNA, TR – translated mRNA, FC – fold change.

The composition of 3’ UTRs can influence both transcript stability and translatability (Mayr & Bartel, 2009; Theil *et al*, 2018; Tushev *et al*, 2018). Considering all editing sites detected in the dataset, 65% occurred in the coding sequence (CDS), while 35% occurred in the 3’ UTR (Figure 4C). The average 3’ UTR length of coding genes quantified in this dataset was only 14.6% of average CDS length, indicating that RNA editing is enriched within this region. To address whether editing of 3’ UTRs influences translation rates, linear regression was performed for CDS-edited and 3’ UTR-edited genes with respect to translational regulation (Figure 4D). Genes with a higher frequency of editing in their 3’ UTRs under DR were more likely to be translationally suppressed (r= - 0.887, p= 1e-5). No significant correlation was observed for genes CDS-edited messages (r= 0.195, p = 0.41). Thus, editing events localized to the 3’ UTR negatively correlate with translation under DR.

Among the genes with differential A-to-I editing in their 3’ UTRs was *eif2-alpha,* a subunit of the eukaryotic translation initiation factor 2. This factor is required for delivering charged tRNA^Met^ to the ribosome to initiate translation (Sokabe *et al*, 2012). Editing of the 3’ UTR of *eif2-alpha* was observed in a comparison of wild type worms to *adr-1* adenosine deaminase mutants (Washburn *et al*, 2014). Two edits were present in the 3’UTR of *eif2-alpha* with significantly different frequencies between total and translated RNA under DR (Figure EV3B and C). In accordance with results for mean editing frequency in Figure EV4A, the editing frequency at both sites was lower in the translated fraction under DR, suggesting that transcripts edited at these loci are translationally suppressed. In line with this, the robust translational promotion of *eif2-alpha* under AL conditions was partially attenuated under DR (DPAR of −0.295, Figure EV3D).

### Annotated RNA-binding sequence motifs are enriched among differentially translated mRNA under DR

RNA-binding proteins (RBPs) and microRNAs (miRNAs) can alter the propensity of a target transcript to be translated. While they differ in their mode of action, both RBPs and miRNAs rely on sequence motifs to recognize mRNA targets. In addition to *cis*-regulatory elements within UTRs of mRNA and corresponding *trans-*acting factors, other characteristics including length and secondary structure can affect translatability (Ringnér & Krogh, 2005; Rogers *et al*, 2011; Zid *et al*, 2009). Therefore, we characterized the 3’ UTRs of promoted and suppressed DPAR genes in terms of length, GC content, free folding energy, and presence of RBP or miRNA motifs to determine their relevance in gene regulation under DR.

The average 3’ UTR length of promoted (148.2 bp) DPAR genes were significantly shorter than that of the quantified gene set (180.6 bp; p=8.71E-9, Wilcoxon rank sum test with continuity correction). No significant differences in mean GC content of translationally promoted or repressed 3’ UTRs were observed. The mean minimal folding energy of promoted 3′ UTRs (−19.1 kcal/mol) were significantly lower than that for the total gene set (−21.5 kcal/mol; 7.19E-05, two-sample Wilcoxon rank sum test with continuity correction). In summary, genes translationally promoted under DR tend to have shorter 3’ UTRs with less secondary structure while repressed genes had no distinguishing features in this region (Table 2).

**Table 2.** Characterization of 3’ UTRs of genes with promoted or repressed polysomal association under DR. Promoted – genes with positive DPAR. Repressed – genes with a negative DPAR, All – all quantifiable genes in the dataset, mean GC – the mean GC content of the UTR sequence, avg BP – the mean sequence base pair length, mean MFE - the mean minimum free energy in kcal/mol. sd – standard deviation, n – the number of sequences available for use in the analysis. Values underlined are significantly (Wilcoxon rank sum test, p-val < 0.05) different compared to all quantifiable genes.

|  | 3' UTR |  |  |
| --- | --- | --- | --- |
|  | Promoted | Repressed | All |
| mean BP | <u>148.2</u> | 170.6 | 180.6 |
| sd BP | 124.2 | 137.9 | 142.1 |
| p-val BP | 8.71E-09 | 0.406 | n/a |
| mean GC | 0.267 | 0.279 | 0.271 |
| sd GC | 0.073 | 0.071 | 0.067 |
| p-val GC | 0.121 | 0.205 | n/a |
| mean MFE | <u>-19.1</u> | -25.1 | -21.5 |
| sd MFE | 27.0 | 32.9 | 25.6 |
| p-val MFE | 7.19E-05 | 0.912 | n/a |
| n | 446 | 81 | 3675 |

To search for sites of potential interaction with RBPs, we used experimentally determined and/or highly conserved motifs for *C. elegans* maintained at the Catalog of Inferred Sequence Binding Preferences of RNA binding proteins (CISBP-RNA) database (Ray *et al*, 2013). Motif enrichment was calculated separately for 3’ UTRs of translationally promoted or suppressed genes relative to the frequency with which motifs occur among all annotated 3’ UTRs in *C. elegans*. 11 RBP motifs were significantly enriched (p < 0.05) in either one or both sets of DPAR genes (Figure 5A). The motifs for *tair-1/2, tair-3*, and *exc-7* were enriched among both promoted and suppressed genes. Target motifs for *mex-5/6*, *unc-75*, *sup-12*, *sup-26*, *etr-1*, *sup-49*, and *asd-2* were enriched almost exclusively among promoted DPAR genes, whereas the target motif for *pab-1* was enriched exclusively among translationally suppressed genes.

**Figure 5.**
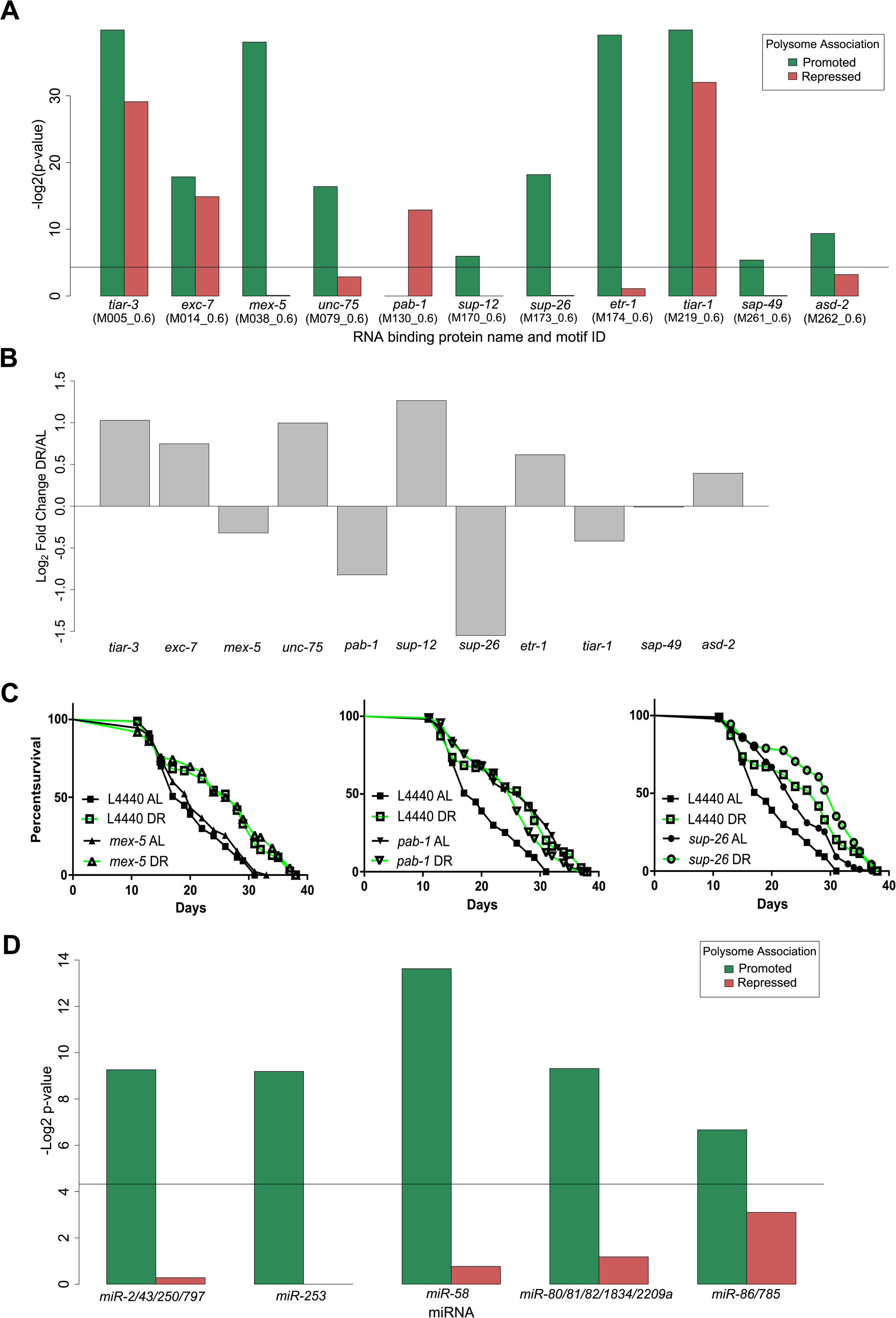
*Trans*-acting factors show enrichment of putative targets among differentially translated genes under DR. **A)** RNA binding protein motifs significantly enriched in 3’ UTRs among differentially translated genes. Black line represents significance threshold of p < 0.05. The CISBP-RNA database motif ID associated with each RNA binding protein is given in parenthesis. **B)** Regulation of RBP genes with enriched motifs under DR. Fold change in the translated fraction is given (* p < 0.05, ** p < 0.01, *** p < 0.001, likelihood test). **C)** RNAi of RBP genes *sup-26* (p=1.39E-04) and *pab-1* (p=2.60E-06), but not *mex-5* (p=0.19), extend median lifespan under well fed conditions compared to controls (n > 80, long-rank test). Representative lifespans are shown, full dataset found in table EV6. AL - ad libitum, DR – Dietary restriction. **D)** miRNA binding sites enriched among translationally promoted or repressed genes under DR. Black line represents significance threshold of p < 0.05. Names of the conserved microRNA families are given.

Of these RBPs, MEX-5/6, SUP-26, and PAB-1 were reported to influence translation of mRNA in *C. elegans*. MEX-5/6 interfere with other RBPs in early differentiation stages during development (Oldenbroek *et al*, 2013). SUP-26 binds the sex determination gene *tra-2* in a 28-nucleotide repeat element located in its 3’ UTR to lower protein expression, most likely by inhibiting translation (Mapes *et al*, 2010). *pab-1* encodes polyA-binding protein, which recent evidence suggests is important for mediating repression and deadenylation through interactions with miRNAs targeting the same transcript (Flamand *et al*, 2016). However, it is primarily considered a pro-translation factor. Each of these RBPs were, themselves, downregulated translationally under DR (Figure 5B). Unlike MEX-5/6 and SUP-26, *PAB-1* targets were enriched among translationally repressed DPAR genes. We tested the requirement of these genes for longevity-and DR-induced longevity using RNAi. Knockdown of *mex-5* had no significant effect on longevity under AL or DR (Figure 5C left, Table EV6). Knockdown of *sup-26* under AL increased median lifespan by 26% (Figure 5C middle, Table EV6). However, knockdown of *sup-26* under DR did not significantly extend lifespan compared to controls on DR. Under AL, knockdown of *pab-1* extended median lifespan by 36% (Figure 5C right, Table EV6) in agreement with previous reports with its effect on longevity (Reis-Rodrigues *et al*, 2012). Under DR, knockdown of *pab-1* was not longer lived than DR controls. Our results for *sup-26* and *pab-1* suggested they are epistatic to DR, as their knockdown under DR didn’t result in an additional increase in lifespan. As *pab-1* encodes for poly A binding protein, a factor that increases translation of mRNA to which it is bound, its downregulation under DR may be pro-longevity as a general effect of widespread decreased translation.

The genes *unc-75* and *asd-2*, which were upregulated under DR, and whose target genes were translationally promoted, both have reported activities in RNA-binding and splicing (Kuroyanagi *et al*, 2013, 2007; Boateng *et al*, 2017; Ohno *et al*, 2012). We used RNAi for *unc-75* and *asd-2* to determine whether their expression was necessary for extended lifespan under DR. If their expression is necessary for DR-induced longevity, then we would expect RNAi treatment under to DR to reduce longevity. Knockdown of *unc-75* under DR increased median lifespan by 11%, suggesting that *unc-75* is not required for DR induced longevity. However, *unc-75* knockdown resulted in a 47% increase in longevity under AL. RNAi treatment of *asd-2* did not significantly reduce longevity under DR compared to controls. Thus, neither *unc-75* nor *asd-2* were necessary for longevity under DR. However, to our knowledge, this is the first study to show *unc-75* is a robust regulator of longevity.

To investigate potential miRNAs that may influence selective translation, we searched for enrichment of miRNA binding sites among translationally promoted and suppressed genes under DR. Predictions of conserved miRNA family targets within 3’ UTRs were taken from targetscanworm.org (Jan *et al*, 2011). Targets of miRNA families miR-2/43/250/797, miR-253, miR-58, miR-80/81/82/1835/2209a, and miR-86/785 were significantly enriched (p < 0.05, Fisher’s exact test) among promoted DPAR genes (Figure 5D). No significant enrichment of targets was found among repressed DPAR genes. The most commonly observed function of miRNAs is silencing gene expression either though degradation of the target or translational suppression (Catalanotto *et al*, 2016). Therefore, if these miRNAs are playing a role in promoting translation under DR, we predict that it is because their expression is downregulated. In line with this, *mir-58* and *mir-80* were shown previously to be downregulated under DR in *C. elegans* (Kogure *et al*, 2017; Vora *et al*, 2013). Deletion of *mir-80* recapitulates many of the physiological and longevity benefits of DR (Vora *et al*, 2013). Furthermore, *mir-58* was recently shown to be partially required for lifespan extension in a genetic model of DR in *C. elegans* (Zhang *et al*, 2018). Based on our *in silico* analysis, the additional miRNA families included in Figure 5D may also play a role in longevity by alleviating suppression of their target mRNAs under DR.

### Intron retention increases under DR

The alternative splicing (AS) of mRNA can produce gene isoforms that are differentially translated. AS can lead to transcripts with premature termination codons (PTC) that are targeted for degradation by nonsense mediated decay (NMD), an mRNA quality control surveillance mechanism that degrades aberrant mRNA. However, the full range of NMD targets extends beyond aberrant gene products and some studies suggest AS and NMD are coupled (AS-NMD) to enable regulation of specific genes via translation (Ge & Porse, 2014; Barberan-Soler *et al*, 2009). Exactly how NMD targets non-aberrant mRNA is unknown, but these and other studies point to the existence of an underappreciated form of gene regulation.

To assess AS-NMD in the current study, we focused on a type of AS in which an intron is retained in the mature mRNA, which usually leads to out of frame translation and one or more PTCs. An average of 1.5% of reads aligned to introns under AL, which was increased to 2.2% under DR (p=0.002, two way t-test, Figure 6A). The change in intron retention under DR suggests either diminished NMD, increased AS, or both. Next, we quantified intron retention in the polysome-associated fraction. NMD takes place during a pioneering round of translation, so we expected mRNAs with retained introns to be depleted among polysomes. Indeed, intron retention was lower in the translated fraction than the total fraction under both AL (1.0%, p-val 0.002) and DR (1.2%, p-val = 0.003). However, the presence of intronic reads in the polysomal fraction suggests that their associated transcripts escaped NMD. Further, while more intron retention was detected in the translated fraction under DR than AL (two way t-test, p = 0.017), the ratio was smaller than that observed between total and translated under AL (p = 0.006, two way ANOVA). Thus, while there are more retained introns among both total and translated mRNA under DR, there is a decrease in the frequency with which intron-bearing mRNA makes it into the translated fraction. These results suggest that intron retention, like 3’ UTR editing, is a method to inhibit protein synthesis of specific genes during adaptation to DR conditions.

**Figure 6.**
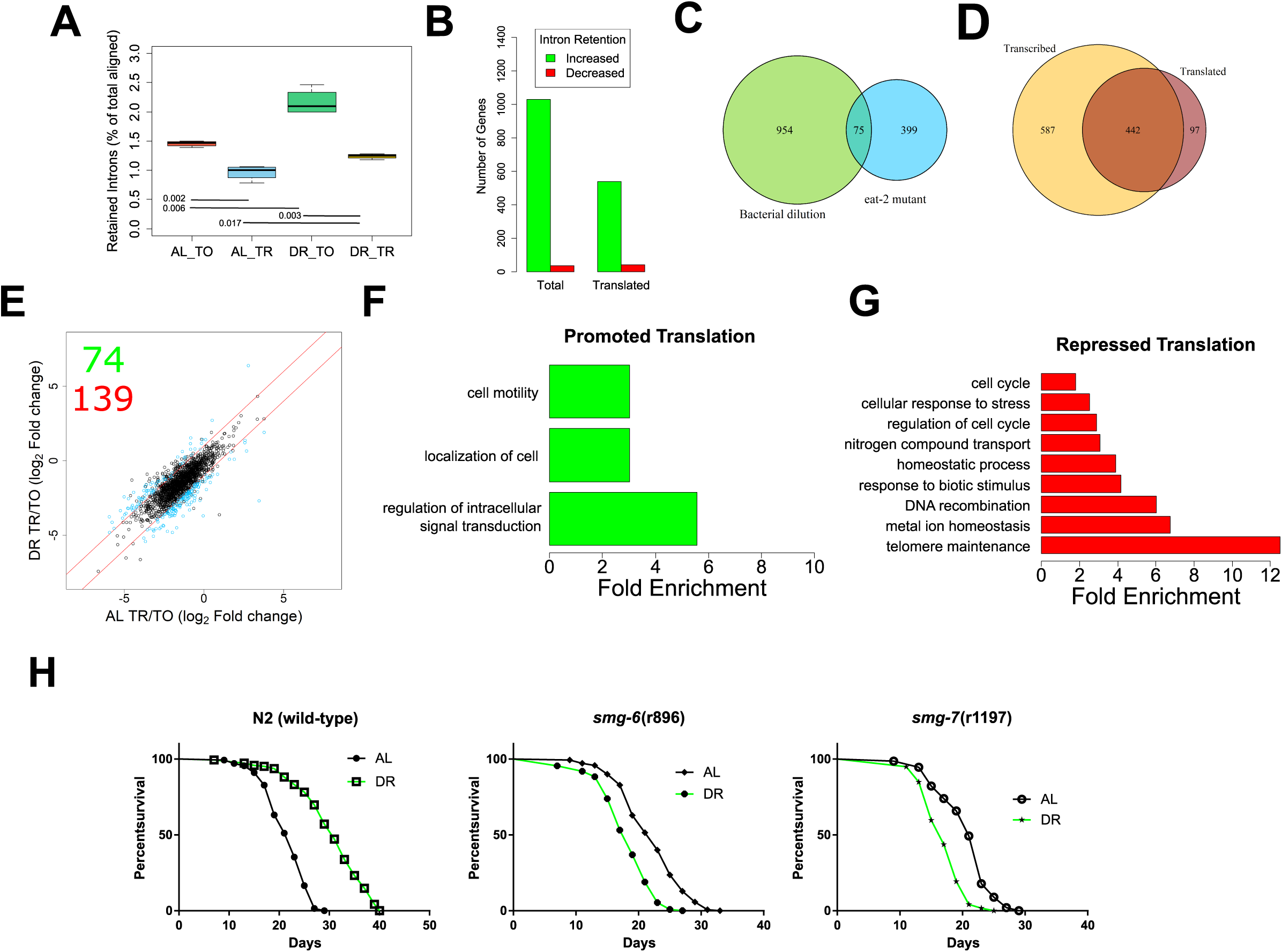
Intron retention is increased under DR to drive changes in translation **A)** The reads aligning to introns as a percent of total aligned. p-values from two-tailed t test, n = 4. **B)** Number of genes with increased or decreased intron retention under DR in either the total or translated fraction. **C)** Venn diagram showing the overlap of genes with intron retention under DR by bacterial dilution in the current study and those identified in the *eat-2* genetic model of DR. **D)** Overlap of genes with increased intron retention under DR in the either the total or the translated fraction. Only genes with > 2 fold change with FDR > 0.05 were considered. **E)** Retained intron transcripts translationally promoted or repressed significantly (> 2 fold, FDR <0.05) under DR. Black circles represent genes with similar translation between diets, blue circles represent genes with two-fold or greater DPAR. Red lines delineate the fold 2 fold change cutoff. Inset, 74 genes were translationally promoted and 139 were translationally repressed. Biological GO terms enriched among retained intron genes with **F)** Promoted or **G)** Repressed translation under DR. p < 0.05, modified Fisher’s Exact test. Functionally similar GO terms from identical groups of genes were manually removed. **H)** Mutants defective in NMD genes *smg-6* or *smg-7* showed a reduction in lifespan under DR, in contrast to wild-type worms. (p< 0.001, long-rank test). Representative lifespans are shown, full dataset found in table EV10. AL - ad libitum, DR – Dietary restriction, TO – total fraction, TR – translated fraction.

Having established that there are dynamic changes in the global frequency of intron retention under DR, we sought to determine changes in the distribution of intron retention on a gene-by-gene basis. Using a fold-change threshold of 2 and a FDR cutoff of 0.05, 1,029 genes showed increased intron retention and 36 exhibited decreased intron retention under DR (Figure 6B). There was a significant overlap in genes with intron retention in the current study with those determined to have intron retention in an *eat-2* genetic model of DR in *C. elegans*, where increased retention of introns was also observed (Figure 6C; Fisher’s exact test, p = 0.01; Tabrez et al. 2017). In the translated fraction, 539 genes showed increased intron retention under DR, while 42 genes had decreased intron retention (Figure 6B). This suggests that changes in intron retention were due to an increase in the number of genes with retained introns. This also suggests that the increased frequency of introns under DR shown in Figure 6A was not simply due to increased frequency of introns within a set of genes common to both AL and DR conditions.

Comparison of genes with increased intron retention under DR in the total and translated fractions shows that most, but not all, of those in the translated pool are a minority subset of genes in the total fraction (Figure 6D). Interestingly, 97 genes exhibited a higher level of intron retention among translated mRNA than among total mRNA, suggesting that the retained introns promoted association with translation machinery in these instances (Figure 6D). Results indicated that intron retention was not always associated with exclusion from translation. This suggested there was more than just nonsense associated with these transcripts and we further investigated the connection between intron-bearing mRNAs and translation.

To identify dynamic changes in translation of intron-bearing genes between AL and DR, we employed DPAR-type analysis similar to that carried out in Figure 2D. 74 intron-retaining genes were associated more with polysomes under DR than AL and 139 were associated less (Figure 6E; Table EV8). Those intron-bearing transcripts translationally promoted under DR were enriched for biological processes related to cell localization and signal transduction, while those translated less under DR and were enriched for biological processes including cell cycle, stress response, and telomere maintenance (Figure 6F-G, Table EV9). Enrichment of genes governing specific biological process suggests that NMD may be selective in a manner that regulates differential gene expression as opposed to just being a quality control process.

While we expect the majority of gene products with retained introns to be non-functional due to frame shift and/or the presence of PTCs, we sought to examine these assumptions empirically and inspect the structure of genes with retained introns. We examined some of the upregulated genes more closely to confirm that intron retention was associated with inclusion of one or more PTCs. Table EV8 shows the frequency of intron retention for individual genes. We first investigated the basic helix-loop-helix protein (bHLH) *mdl-1* due to its role in regulating *C. elegans* lifespan (Nakamura *et al*, 2016) and because it was previously reported to undergo intron retention (Tabrez *et al*, 2017). The predicted CDS of the intron bearing isoform of mdl-1 contained an out of frame PTC occurring after its annotated bHLH domain (Figure EV4A). Therefore, while the retained intron isoform would encode for a truncated protein it could still have a function though the expressed bHLH domain.

The gene F13G3.6 was investigated as it was reported to have a PTC bearing isoform whose expression is dependent on the activity of the NMD gene *smg-2* in *C. elegans* (Son *et al*, 2017). This gene is similar to human B3GNTL1, which plays a role in elongating N-glycans on mucin proteins. Exons upstream of the intron contained a Glycosyl transferase family 2 domain. Interestingly, the retained intron results in a single PTC with no frame shift (Figure EV4A). In such instances, it is possible that translational read-through of the PTC occurs to produce a full length product (Oren *et al*, 2014).

Last, we examined the selenoprotein *trxr-1* as it mediates healthy longevity in response to treatment with the organic form of selenium (Chang *et al*, 2017) and only retained a portion of the annotated intron. The retention event resulted in the addition of 6 base pairs to the 5’ end of exon 5, which did not induce a PTC or frame shift (Figure EV4C). Therefore, while unlikely to be directly regulated by NMD, this isoform may represent a new, previously annotated isoform of *trxr-1* that is better translated under DR.

### NMD genes *smg-6* and *smg-7* are required for DR-mediated lifespan extension

In *C. elegans*, NMD is governed by the activity of several genes, including *smg-1/2/3/4/5/6/7*, which are responsible for the detection of aberrant transcripts, releasing the stalled ribosome, and degrading the transcript (Hug *et al*, 2016). Given that the translation of NMD prone transcripts appeared to be regulated by DR and that NMD has been shown to play a role in ILS mediated longevity (Son *et al*, 2017), we hypothesized that NMD plays a role in DR induced longevity. We tested the importance of NMD in longevity using mutants defective in *smg-1/2/3/4/5/6/7* subject to AL or DR starting at adulthood. Interestingly, the lines *smg-6(r896)* and *smg-7(r1197)* were shorter lived under DR than AL (p-val < 0.0001 for both; Log-rank test), demonstrating their requirement for DR-induced longevity (Figure 6H). In comparison, the other *smg* mutants (*smg-1/2/3/4/5*) were longer lived under DR than AL, albeit this extension was attenuated compared to wild-type worms (Table EV10). The genes *smg-6* and *smg-7* are each thought to participate in NMD by degrading the transcript after it has been detected and the ribosome released. While *smg-6* is thought to degrade transcripts directly by its endonuclease activity, *smg-7* is thought to degrade transcripts by recruiting deadenylation factors. Together, these observations raise the possibility that the selective degradation of NMD prone transcripts by *smg-6* and *smg-7* can dictate which transcripts are translated under DR to influence longevity.

## Discussion

Translation is expensive with respect to the energy and raw materials that it consumes. Low nutrient levels lead to low global translation. However, our understanding of how the stoichiometry of protein synthesis for specific individual gene products changes under DR through selective translation is limited. To elucidate this process, we sought to identify translationally regulated genes as well as potential mechanisms of post-transcriptional regulation that influence their translation. Utilizing a combination of polysome profiling and mRNA sequencing, we generated diagnostics of the transcriptome and translateome. *In silico* analysis was used to assess the relationship between these diagnostics and how this relationship is changed when nutrients are restricted. It also provided a means by which to probe differentially expressed genes for annotated *cis*-elements and corresponding *trans*-acting factors, generating a roadmap for future analyses testing mechanisms of post-transcriptional regulation that contribute to differential expression. The use of RNAi allowed us to determine which differentially expressed genes and cellular processes, like NMD, are important contributors to the increased lifespan associated with DR.

An important early step in trying to understand how somatic protection is conferred by a health-promoting intervention like DR is to understand how expression of genes change to meet the requirements of adaptation. Three genes among the 100 showing the greatest transcript-level decrease in expression under DR increased lifespan when knocked-down with RNAi under AL. *fbxb-24* and *sdz-*33 encode highly related F-box containing proteins. The F-box motif was first observed as a constituent of SCF ubiquitin-ligase complexes and helped confer specificity to proteolytic substrates (Skowyra *et al*, 1997). Since then, many other genes encoding F-box motifs have been identified. The motif is structured with about 50 amino acids that function in protein-protein interactions, which are highly abundant in *C. elegans* (Kipreos & Pagano, 2000). The third gene was D1086.3, a gene of unknown function that was also the only one tested for lifespan that was also selectively downregulated via translation.

Selective translation of stress responsive genes important for longevity has been previously described in *C. elegans* ILS mutants (McColl *et al*, 2010) and in a model of restricted nutrient-responsive translation (Rogers *et al*, 2011). In *Drosophila melanogaster*, selective translation enhanced synthesis of genes encoding mitochondrial components on a form of DR involving restriction of yeast extract (a primary source of protein; Zid et al. 2009). Inhibition of the nutrient-responsive TOR complex in mouse embryonic fibroblasts selectively suppressed ribosomal genes and promoted translation of several transcription factors, including *Foxo1* (Thoreen *et al*, 2012). We observed translational suppression of ribosomal genes in the current study as well as other genes involved in translation (Figure 2G, Table EV3). Two of these, *iff-1* and *pab-1*, are translation initiation factors that increased lifespan when knocked down in fully fed wild type animals.

In addition to direct regulators of translation, other repressed DPAR genes involved in lifespan regulation encode chaperones. The largest increase in median lifespan was observed for *daf-21*, the *C. elegans* ortholog of human HSP90. Although it has been previously reported to increase lifespan (Sutphin *et al*, 2017; Horikawa *et al*, 2015), this is the first study to demonstrate its translational regulation in the context of DR. DAF-21/HSP90 serves to promote growth and reproduction by keeping growth-related kinases and transcription factors in a near-native conformation that facilitates their activation (Mitra *et al*, 2016). HSP90 also sequesters the HSF-1 heat shock transcription factor, inhibiting its ability to trimerize and upregulate expression of heat shock protein genes (Zou *et al*, 1998). The translational downregulation of *daf-21* under DR makes sense given that there should be less need for activation of growth related kinases and transcription factors in low energy conditions when nutrients are depleted. The chaperonin gene, *cct-4*, was translationally down regulated under DR and increased lifespan upon knockdown despite the fact that loss of *cct-4* is associated with a reduction *skn-1* transcriptional activity (Kahn *et al*, 2008). *skn-1* is the only worm ortholog of mammalian nuclear factor-erythroid-related factor (Nrf) transcription factors and functions in the p38 MAPK pathway to regulate the oxidative stress response. Its transcriptional activity is inhibited by insulin-like signaling (Tullet *et al*, 2008) and it is considered to be a longevity factor by promoting somatic maintenance. It is unclear why low *cct-4* translation supports increased lifespan given its positive correlation with at least one *skn-1* target, *gst-4*. It is possible that other *skn-1* targets are not similarly affected by *cct-4* expression.

Translationally downregulated genes D1086.3 and *kbp-1* were, for the first time, identified as lifespan regulators in our RNAi screen. Interestingly, we did not observe lifespan extension in our RNAi screen for certain repressed DPAR genes previously shown to regulate lifespan. For example, expression of *skr-1* and *ncbp-2* are positively associated with longevity, while expression of *cyc-2.1* is negatively associated with longevity but had no effect when knocked down using RNAi in the present study. The importance of *skr-1* for increased lifespan is specific to attenuated ILS in *daf-2* mutant backgrounds of multiple alleles tested and does not regulate lifespan in an *eat-2(ad1116)* genetic model of DR (Ghazi *et al*, 2007). RNAi targeting the nuclear cap binding protein gene *ncbp-2* was only shown to shorten lifespan when initiated during development, which can also result in severe defects and lethality (Büssing *et al*, 2010). *cyc-2.1* encodes a component of the electron transport chain, which only extends lifespan in N2 wild-type animals when it is knocked down during development. Thus, it is not surprising that these genes did not regulate lifespan in our longevity screen that began during adulthood.

### A novel association between RNA editing and translation under DR

Although RNA editing frequencies were previously shown to be regulated by neuronal stimulation (Tariq *et al*, 2013), regeneration (Witman *et al*, 2013), and hypoxia (Ben-Zvi *et al*, 2013), this is the first study to reveal widespread changes in editing due to nutrient limitation. A-to-I RNA editing increased under DR, especially within the 3’ UTRs, and such editing was associated with translational suppression. Conversely, although editing events occurred in the CDS, they were less frequent and were not associated with translational suppression. Together, these results indicate that RNA editing events are dynamic and likely to play a role in the adaptive response to DR.

While a link between editing of individual genes and longevity remains to be established, polymorphisms in mRNA editing genes have been associated with longevity in humans (Sebastiani *et al*, 2009). The same study also demonstrated that loss the orthologous editing genes in *C. elegans* were required for longevity (Sebastiani *et al*, 2009). This raises the possibility that mRNA editing regulates the selective translation of transcripts with inputs to longevity. A provocative example of this in the current dataset was the editing of the 3’ UTR of *eif-2alpha.* Furthermore, the enrichment of RNA editing sites within 3’ UTRs was shown to overlap frequently with miRNA binding sites (Gu *et al*, 2012). Thus, RNA editing may influence expression by introducing or removing miRNA binding sites.

### RBPs and translation regulation under DR

Our screening of RBP motifs implicated *pab-1* in the regulation of translationally promoted genes under DR. This gene encodes for the cytoplasmic poly-A-binding protein, referred to as PABPC1 in mammals. It has several molecular functions associated with it, the most prominent of which is its ability to bind to poly-adenosine stretches in the 3’ UTR of mRNA and then to associate with factors in the 5’ UTR to promote translation initiation. PABPC1 also regulates the expression of specific mRNA by binding to regions outside of the polyA tail (Smith *et al*, 2014). This mRNA specific regulation can be mediated by interactions with miRNA (Flamand *et al*, 2016), participation in NMD (Brook & Gray, 2012), or through association with the 3’ UTR instead of the polyA tail (Burgess & Gray, 2010). Thus, *pab-1* may be important for multiple forms of differential translation under DR.

### Translational preference for *trans*-spliced products is reversed for the subset of SL genes in operons during adaptation to DR

5’ UTRs of *C. elegans* transcripts are frequently *trans-*spliced, which is a feature not only common to nematodes, but also to flatworms, protozoa, sponges, cnidarians, chaetognaths, crustaceans, rotifers, and tunicates (Pouchkina-Stantcheva & Tunnacliffe, 2005; Zhang *et al*, 2007; Zayas *et al*, 2005; Hastings, 2005; Marlétaz *et al*, 2008). SL sequences in nematodes bear trimethylguanosine caps (m_2,2,7_G; Liou and Blumenthal 1990), in contrast to native 5’ UTRs, which contain a monomethylguanosine (m_7_G) cap (Sonenberg, 2000). Under well fed conditions, we found that *trans*-spliced mRNA was translated better than mRNA with the native 5’ UTR, supporting previous analysis (Yang *et al*, 2017). While this pattern was maintained under DR, it was modestly but significantly attenuated for SL1 spliced genes (Figure 3B).

Interestingly, despite the preference for SL-containing mRNA to be translated under normal conditions, both SL1 and SL2 genes were underrepresented among promoted DPAR genes (Figure 3C, D). This was also true for polycistronic genes contained in operons, which represent a subpopulation of *trans*-spliced products that accounts for ~17% of all protein coding genes in *C. elegans* (Blumenthal & Gleason, 2003). Unlike bacterial operons, nematode polycistrons are separated post-transcriptionally and translated separately. The first gene within an operon is spliced to SL1 and subsequent genes are spliced to SL2. Interestingly, nearly half of the DPAR genes downregulated in the transition from AL to DR conditions reside in operons. Although it is not clear how they are downregulated, co-regulation of multiple genes by a single promoter could represent an optimization of gene expression for growth and reproduction under favorable conditions. Unfavorable conditions may then necessitate differential expression of genes under the same promoter, which could be achieved through translational regulation. However, due to the differences in the broader trend for translational regulation of trans-spliced genes from the subset of those in operons, we surmise that the presence of SLs, alone, cannot account for the differential translation of operonic genes under DR.

### DR increases AS products and differentially regulates NMD targets

We found evidence that DR increases the prevalence of AS and alters NMD. Furthermore, we found that certain NMD genes that play a role in target specificity and degradation are required for the benefits of DR. How AS-NMD is integrated to influence differential gene expression remains to be fully elucidated. With respect to splicing, there is a limited but growing body of evidence pointing to changes in AS according to dietary and metabolic conditions. For example, DR is associated with changes in AS for individual metabolic genes in mammals (Kaminska *et al*, 2014; Gallardo *et al*, 2005; Fernández *et al*, 2009). Conversely, obesity is associated with diminished expression of several splicing factors in liver and skeletal muscle in a manner that contributes to increased lipogenesis (Pihlajamäki *et al*, 2011). In a recent non-DR related study of splicing in the tea plant *Camillia sinensis*, the predominant form of AS was intron retention, which occurred in genes related to the spliceosome and mRNA processing (Zhu *et al*, 2018). That regulators of splicing are, themselves, highly subject to alternative processing is conserved in animal studies. In mammals, AS is regulated by members of the SR protein family that alternatively splice a portion of their own transcripts (Lareau *et al*, 2007). These splice variants are normally targeted by NMD to limit total SR protein expression. Our analysis of intron retention suggested that coupling of AS-NMD is modulated under DR, in part, by changing the selectivity towards which transcripts are detected and degraded. Observations are in line with those obtained from cell culture exposed to rapamycin, a drug that inhibits the DR-related TOR pathway, in which the selectivity of NMD towards transcripts was also modulated (Martinez-Nunez *et al*, 2016). Taken together, studies point to an adaptive mechanism of DR that functions by controlling productive splicing and turnover.

The idea of regulated AS-NMD was previously invoked to explain differential expression during development in *C. elegans* (Barberan-Soler *et al*, 2009). Recently, (Tabrez *et al*, 2017) used a genetic model of DR in *C. elegans* and identified over 400 differential intron retention events. The authors suggested that alternative splicing under DR is needed to diversify the proteome in a way that is conducive for longevity and additionally requires NMD to fine tune gene expression. The same mutant was found to have increased splicing in the early, healthy part of life (Heintz *et al*, 2016). Thus, while it is clear that AS and intron retention is modulated by DR in a way that may affect longevity, the roles of AS-NMD on differential gene expression are, as yet, unclear.

This study was designed to characterize the post-transcriptional landscape of gene expression in response to DR and then survey the mechanisms which may govern this regulation. One limitation to this approach was the use of whole worms as opposed to specific cells lines or tissues. While the use of whole worms allows for the quantification systemic changes on translational regulation or effects occurring in large tissues like the intestine, tissue specific effects, like RNA editing in the neurons, are likely lost by averaging the effect over all other tissues. The use of tagged RNA binding proteins, like poly A binding protein, expressed by tissue specific promoters has been used in *C. elegans* to quantify tissue-specific gene expression in total mRNA pools (Blazie *et al*, 2015, 2017). Future studies may rely on similar technology to identify genome-wide tissue-specific patterns of differential processing and expression.

## Materials and Methods

### Polysome Profiling

Polysome profiling was performed as previously described (Pan *et al*, 2007). Day 4 adult worms were used for polysomal profiles. 100 ul of pelleted worms were homogenized on ice in 350 ul of solubilization buffer (300 mM NaCl, 50 mM Tris HCl [pH 8.0], 10 mM MgCl2, 1 mM EGTA, 200 g heparin/ml, 400 U RNAsin/ml, 1.0 mM phenylmethylsulfonyl fluoride, 0.2 mg cycloheximide/ml, 1% Triton X-100, 0.1% sodium deoxycholate). After homogenation, an additional 700 ul of solubilization buffer was added and the sample was incubated on ice for 30 minutes. Debris was pelleted by centrifugation at 20,000 g for 15 min at 4°C. Of the resulting supernatant 900 ul was applied to the top of a 10-50% sucrose gradient in high salt resolving buffer (140 mM NaCl, 25 mM Tris-HCl [pH 8.0], 10 mM MgCl2) and centrifuged in a Beckman SW41Ti rotor at 38,000 rpm for 90 min at 4°C. Gradients were fractionated using a Teledyne density gradient fractionator with continuous monitoring of absorbance at 252 nm. RNA was extracted from ploysome fractions and unfractionated samples using Trizol LS reagent according to the manufacturers protocol (Invitrogen Corp., Carlsbad, CA, USA). Eukaryote total RNA Nano chips were used in an Agilent Bioanalyzer for quality control of the resulting RNA.

### Alignment of reads to the *C. elegans* genome

Alignment of RNA-seq reads from each biological replicate to *C. elegans* genome version WS220 was performed using the splice junction mapper TopHat (ver 2.0.8.b) (Kim *et al*, 2013). TopHat was supplied with transcript annotations from ENSEMBL version 66 (Flicek *et al*, 2011), so that preference of aligning reads is given to the transcriptome first (remaining reads are converted to genomic mappings and merged with the final output). Default settings of Tophat were used with the following exceptions: the mean mate inner distance was set to 200 bp, microexon search was enabled, and the library type was set to ‘fr-unstranded’ to indicate that unstranded paired end reads were used.

### Differential Expression

Aligned reads were counted per gene using the python script HTseq (Anders *et al*, 2015). Differential expression and dataset normalization was performed using the Bioconductor package edgeR (Robinson *et al*, 2010). Normalization in edgeR adjusted for RNA composition to ensure that highly expressed genes which consume a large portion of the RNA pool didn’t result in the under-sampling of other genes. Additionally, as our data set contained multiple factors, dispersion of the gene counts were estimated tagwise using the Cox-Reid profile-adjusted likelihood method (Cox & Reid, 1987). Only genes with average counts per million (CPM) of eight or greater across all conditions were considered reliably quantifiable for differential expression. A threshold of eight CPM was chosen as it was the point where the trend in the biological coefficient of variation approached the common variation. Differential expression was calculated pairwise between groups using a general linear model and the resulting p-values were adjusted for multiple testing using the Benjamini-Hochberg method (Benjamini & Hochberg, 1995). The Differential Polysome Association Ratio (DPAR) was calculated using the following GLM using EdgeR’s makecontrast() command as follows: (DR_TR - DR_TO)-(AL_TR - AL_TO).

### Expression Profile Clustering

Genes with a significant two-fold or greater difference in relative expression between total and polysome-associated mRNA under DR but not AL were clustered together. Prior to clustering, the expression of each gene among the four experimental groups in was centered around its mean expression in the dataset. Clustering was performed with the Partitioning around Medoids (PAM) algorithm using the pam() function and evaluated using silhouettes widths (Rousseeuw, 1987) with the silhouette() function, both contained in the R package ‘Cluster’ (Maechler *et al*, 2013). The maximal number of clusters to use was determined by reiteratively running the PAM algorithm while sequentially increasing the number of clusters. Based on the experiences of Kaufmann and Rousseeuw (Kaufman & Rousseeuw, 2005), a silhouette width cuff-off of 0.25 was used as the lower limit for a cluster to have substantial structure. The highest cluster number among all iterations with all clusters having substantial structure was three.

### RNA-binding protein motif enrichment

Experimentally determined and homology inferred position weighted matrices for RNA-binding motifs of *C. elegans* proteins were downloaded from the CISBP-RNA database (Ray *et al*, 2013). Enrichment for each motif among genes translationally promoted or suppressed under DR was calculated using the R package PWMEnrich (Stojnic & Diez, 2013). Background correction for nucleotide composition was applied with the function makePWMLognBackground() using 3’UTRs from all quantifiable genes. 3’UTRs were exported from the *C. elegans* genome version WS220, 3’UTRs shorter than 12 bp were excluded from the analysis.

### miRNA motif enrichment

Predicted targets of conserved miRNA families among nematodes were imported from http://www.targetscan.org/ release 6.2 (Jan *et al*, 2011). Only targets with a probability of conserved targeting (P_ct_) > 0.8 were considered. Enrichment of miRNA binding sites among translationally promoted or suppressed genes compared to all quantifiable coding genes was calculated using Fishers exact test.

### Spliced leaders

Annotations of the predominant spliced leader present in *C. elegans* transcripts were taken from (Allen *et al*, 2011) supplement table 2. Unannotated genes were considered not to undergo *trans-*splicing and have their native 5’UTRs. Comparison of translational regulation between AL and DR of genes with native, SL1, or SL2 5’UTRs was performed using the Wilcoxon rank sum test with continuity correction.

### Intron retention

Reads aligning to intronic or exonic sequences were counted for each biological replicate using HTseq (Anders *et al*, 2015) based on transcript annotations from ENSEMBL version 66. A minimum of four reads aligning to introns within a gene was required to include in the analysis. The amount of global intron retention was expressed as the percent of reads aligning to introns over all aligned reads. Significant differences in global intron retention were determined using Student’s T-test. To calculate intron retention occurring between diets in the total or translated fraction for individual genes the Wilcoxon rank sum test was used. To detect genes with intron retention differing between RNA fractions in a way dependent on DR, a linear model was used to determine significant interaction effects of diet and RNA fraction on intron retention. This was achieved using the lm() function in R to model the interaction effect of diet and RNA fraction on intron retention for each gene. Multiple testing was corrected for using the Benjamini & Hochberg method.

### Gene ontology term enrichment

Enrichment of biological terms among gene lists was performed using DAVID (version 6.7) (Huang *et al*, 2008, 2009). The category GOTERM_BP_FAT was used for annotations and the list of quantifiable coding genes in our dataset was used as the background. For visualization, similar enriched GO terms based on the same subset of genes were manually removed to reduce redundancy, supplemental tables contain the full results.

### Statistical Analysis

Spearman correlations between gene abundance in the total and translated fractions was calculated using the rcorr() function in the R package Hmisc (Jr *et al*, 2013). The distribution of fold change in gene abundance between the total and translated fractions was compared between AL and DR conditions using a two-sample Kolgomarov-Smirnov test.

### Lifespans

All worm cultures were maintained at 20°C. Worms were subjected to RNAi treatments starting at day one of adulthood to avoid potential effects on development. To prevent contamination by progeny, FuDR was used at a concentration of 0.5 mg/ml in the plate. Worms were scored every other day for survival. Worms were scored as alive if movement was evident after gentle prodding of tail and head with a platinum worm pick. Worms with vulvar bursts or that had crawled off media were censored. Survival was plotted as Kaplan-Meier survival curves and analyzed by log-rank test in GraphPad Prism (version 5).

### RNA editing sites

To search for potential adenosine to inosine RNA editing sites, mRNA-seq reads from each biological replicate were compared to the reference genome WS220 for the presence A to G SNPs using REDItools (Picardi & Pesole, 2013). The last three bases from each were trimmed, and only uniquely mapped reads from properly paired reads were considered. Additionally, for each biological replicate, a minimum of five reads supporting the variation and an editing frequency greater than 10% was required. Significant differences in editing frequencies between experimental groups were determined using the Wilcoxon signed-rank test.

### Gene Set Enrichment Analysis (GSEA)

Comparison of differentially expressed transcripts in the total fraction of our dataset to those detected in a microarray experiment published by Honjoh et al (Honjoh *et al*, 2009) was performed using gene set enrichment analysis version 2.0.9 (Subramanian *et al*, 2005). The processed data from the manuscript was downloaded from Array Express under the genome accession number GSE9682. The processed data were reformatted for GSEA and Affy IDs were replaced with the gene ID for their matching transcript using Ensembl’s Biomart function. Of the 22626 affy IDs, 21459 had a gene name assigned to them; those Affy IDs without a corresponding Gene ID were not included in the analysis. The GSEA was run with 10000 permutations based on shuffling the gene set, using the ‘weighted’ enrichment statistic and ‘signal to noise’ metric for ranking genes.

### 5′ and 3′ UTR characterization

5′ and 3′ UTRs were characterized separately based on length, GC content, and minimum free folding energy. Three groups of genes were considered, genes translationally promoted, genes translationally suppressed, and all quantifiable genes. Gene sequences from the 5′ or 3′ ends were exported from ENSEMBL version 66 using the getSequence() function in the biomaRt R package. As genes often encode for multiple isoforms, 3’ and 5’UTR sequences from the predominately expressed isoforms across the experiment was used for analysis. Quantification of isoform expression was performed using cuffdiff with default parameters (version 2.1.1) and the predominantly expressed isoform was defined as the one with the highest average FPKM in the dataset. Fractional GC content and sequence length was calculated using the alphabetFrequency() function from the biostrings package. The minimum free folding energy was calculated using the RNAfold function of the Vienna RNA package (version 2.4.8) with default settings. The characterized values for translationally promoted and suppressed genes were compared to all quantifiable genes using the Wilcoxon rank-sum test.

## Supporting information

Extended View Figures 1-4

## Data Availability

The raw and processed high throughput sequence data from this publication have been deposited to the Gene Expression Omnibus database https://www.ncbi.nlm.nih.gov/geo/ under accession GSE119485.

### Acknowledgements

The authors would like to thank Ben King, Ph.D. and Joel Graber, Ph.D. for bioinformatics advice as well as George Sutphin, Ph.D. for constructive criticism of the manuscript. This work was supported by grants from the National Institute on Aging of the National Institutes of Health (R21AG056743) and by the Ellison Medical Foundation (AG-NS-1087-13), both to A.R. Additionally, this project was supported by Institutional Development Awards (IDeA) from the National Institute of General Medical Sciences of the National Institutes of Health (grant numbers P20GM0103423 and P20GM104318, respectively).

## Author Contributions

A.R. and P.K. devised the study. A.R. and J.R. carried out experiments and prepared the manuscript. P.K. helped with scientific interpretation. S.S. performed additional experiments.

---

**Figure EV1.**
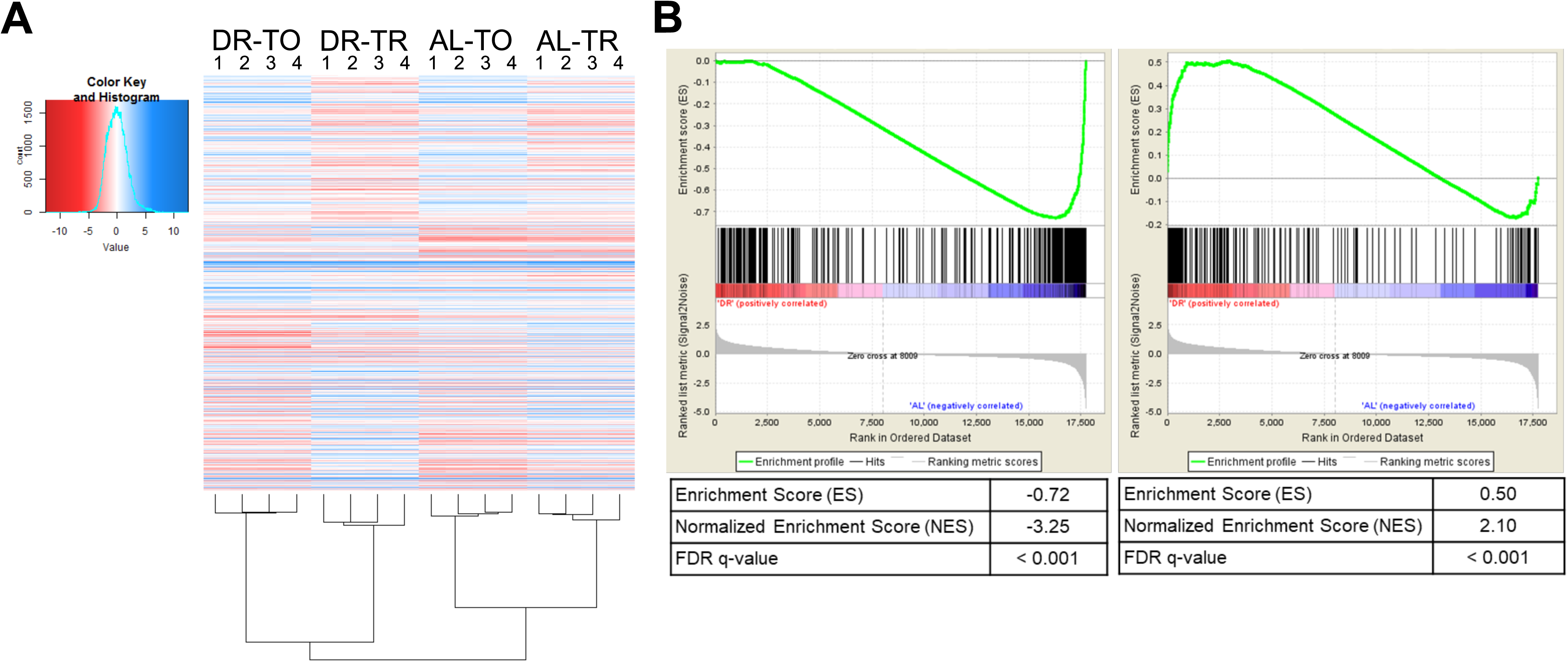
Extended analysis of differential expression datasets. **A)** Heatmap of all quantifiable genes in the dataset demonstrates good clustering by biological replicates. DR – dietary restriction, AL – ad libitum, TO – total mRNA, TR – translated mRNA. **B)** Gene Set Enrichment Analysis comparing the list of genes down-regulated (left) or up-regulated (right) transcriptionally under DR two-fold or more in the present study against the ordered gene set regulated due to intermittent fasting (Honjoh *et al*, 2009). The top enrichment profile shows the cumulative enrichment score as each ranked gene in the list is compared to the gene set. Where each gene in the list falls in the gene set is represented by black bars in the middle section. The bottom of the plot depicts the ranked list metric which measures a gene’s correlation with the treatment. Both lists were significantly (FDR < .001) enriched as compared to the microarray. Better agreement to intermittent fasting was found among the down regulated genes (normalized enrichment score = −3.25) than up-regulated genes (normalized enrichment score = −2.10). More details available in the Materials and Methods.

**Figure EV2.**
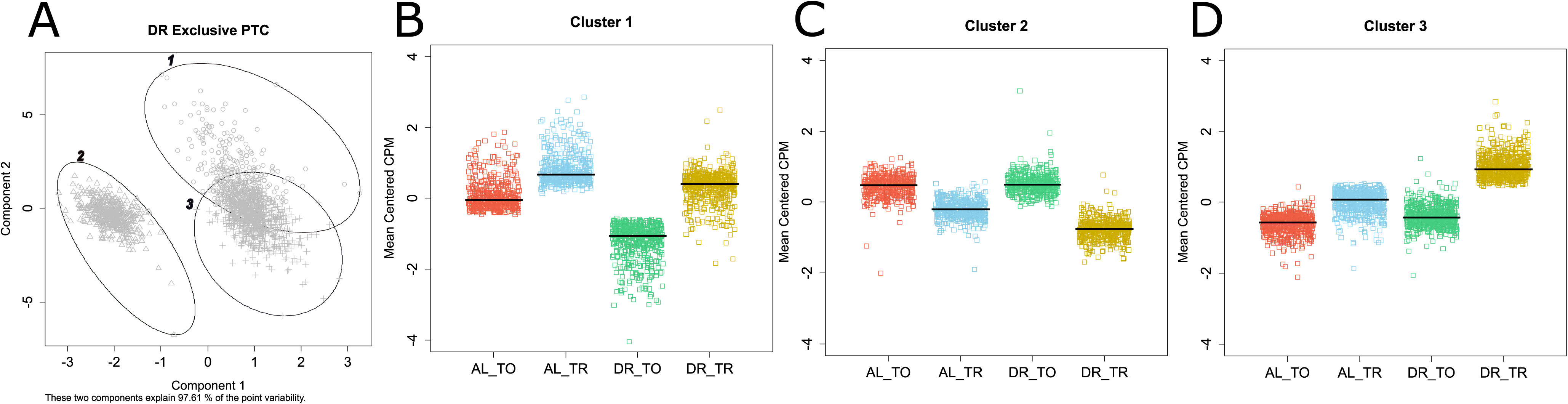
Predominant modes of regulation under DR from unbiased clustering of genes that only exhibit translational regulation under DR. **A)** Genes selectively translated under DR were optimally clustered into three groups and visualized in a 2-dimensional principle component analysis plot representing 97.6% of the variation in the dataset. **B-D)** The mean centered expression of genes in clusters 1 through 3, respectively. Bars represent the median expression of each group.

**Figure EV3.**
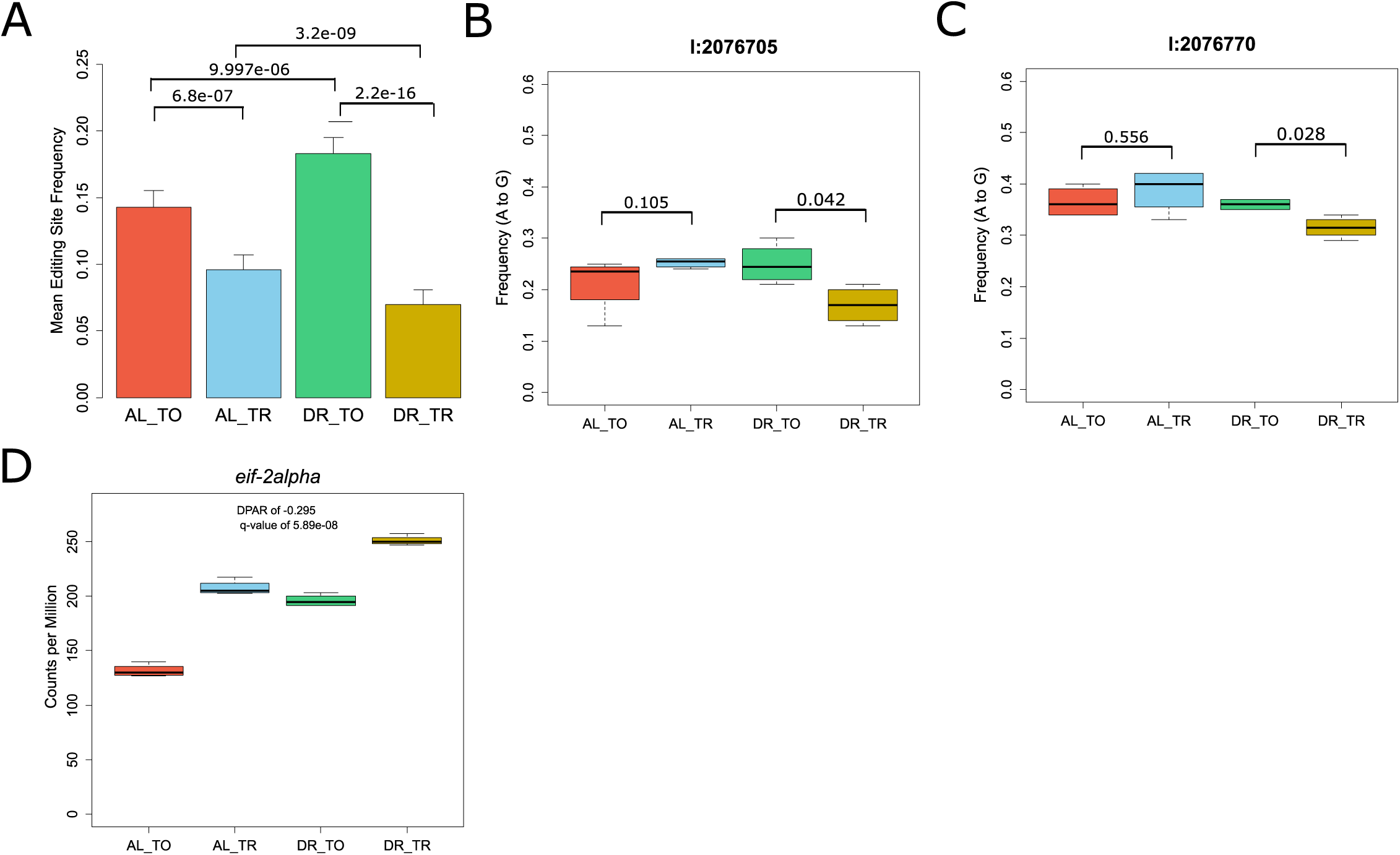
Altered A-to-I editing in 3’ UTR of *eif-2alpha*. **A)** Average frequency of A-to-I editing in total and translated fractions across diets among all 331 sites detected. As with the number of overall sites in Fig 4A, the frequency of edits increases among total mRNA under DR and but is further attenuated within the translated fraction. P-values from Wilcoxon rank sum test between indicated comparisons are given, n=331. Error bars are s.e.m. The mean editing frequency at position **B)** 2076705 bp and **C)** 2076770 bp on chromosome 1 within the 3’ UTR of *eif-2alpha*. These edits decrease in frequency in the translation fraction under DR. p-values are given for comparisons between fractions, Wilcoxon rank sum test, n=4). **D)** Expression of *eif-2alpha* under the total and translated fractions across diets as expressed in the number of reads aligned per million reads generated. TO – total mRNA, TR – translated mRNA.

**Figure EV4.**
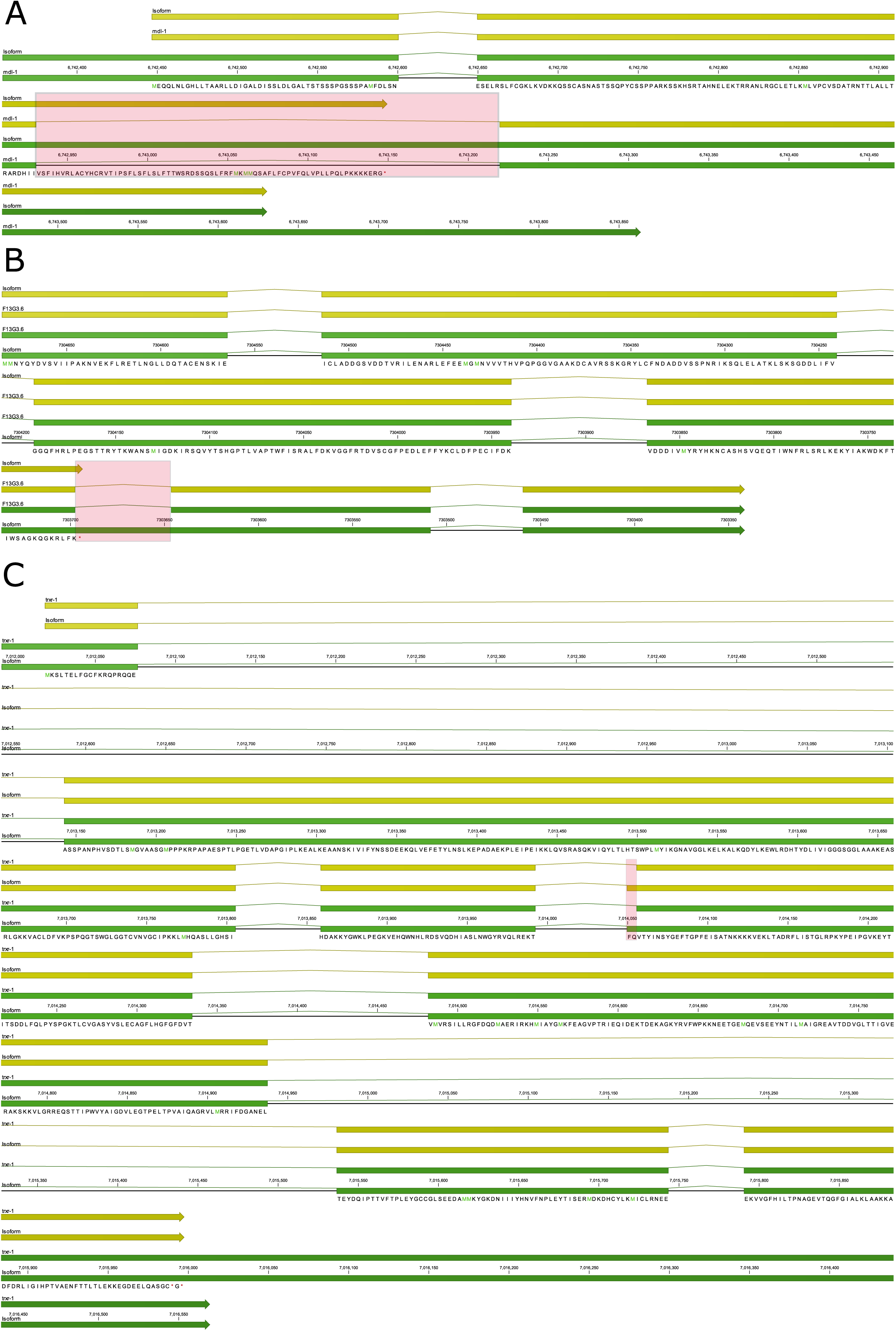
Retained introns change the coding potential of longevity genes *mdl-1*, *F13G3.6*, and *trxr-1*. The mRNA (green features) and predicted coding sequence (yellow features) are shown for the genes **A)** *mdl-1*, **B)** *F13G3.6*, and **C)** *trxr-1* and their isoforms with retained introns that were translationally promoted under DR. Red boxes highlight the differences between the isoforms. The predicted translated sequence is given for the novel isoforms.

## Expanded View table legends

**Table EV1.** List of quantifiable genes significantly (FDR < 0.05) up-or downregulated two-fold or more under dietary restriction in the total mRNA. Explanation of column names are given in the key.

**Table EV2.** Biological process gene ontology terms enriched among genes significantly up-or downregulated dietary restriction in the total mRNA. Terms enriched among genes regulated in common between DR by bacterial dilution in the present study and DR by fasting are given as well are terms enriched among genes that were uniquely regulated by bacterial dilution. Explanation of column names are given in the key.

**Table EV3.** List of quantifiable genes translationally promoted or repressed two fold or more under dietary restriction. Explanation of column names are given in the key.

**Table EV4.** Biological process gene ontology terms enriched among genes translationally promoted or repressed under dietary restriction. Explanation of column names are given in the key.

**Table EV5.** List of translationally repressed and transcriptionally downregulated genes tested in RNAi longevity screen. Wormbase sequence IDs or gene name is given.

**Table EV6.** Summary table of lifespans conducted on using RNAi for RNA binding protein genes. Explanation of column names are given in the key.

**Table EV7.** Frequency of adenosine to inosine edited sites among total and translated mRNA under ad libitum or dietary restriction in C elegans. Explanation of column names are given in the key.

**Table EV8.** List of genes with genes with intron retention levels that occur between total and translated fractions in a way dependent on dietary restriction. Explanation of column names are given in the key.

**Table EV9.** Biological process gene ontology terms enriched among retained intron genes with promoted or repressed translation under DR. Explanation of column names are given in the key.

**Table EV10.** Summary table of lifespans conducted using *smg*-*1/2/3/4/5/6/7* mutants under ad libitum or dietary restriction. Explanation of column names are given in the key.

