## Extended View Figures 1-4 for "Dietary restriction induces post-transcriptional regulation of longevity genes"

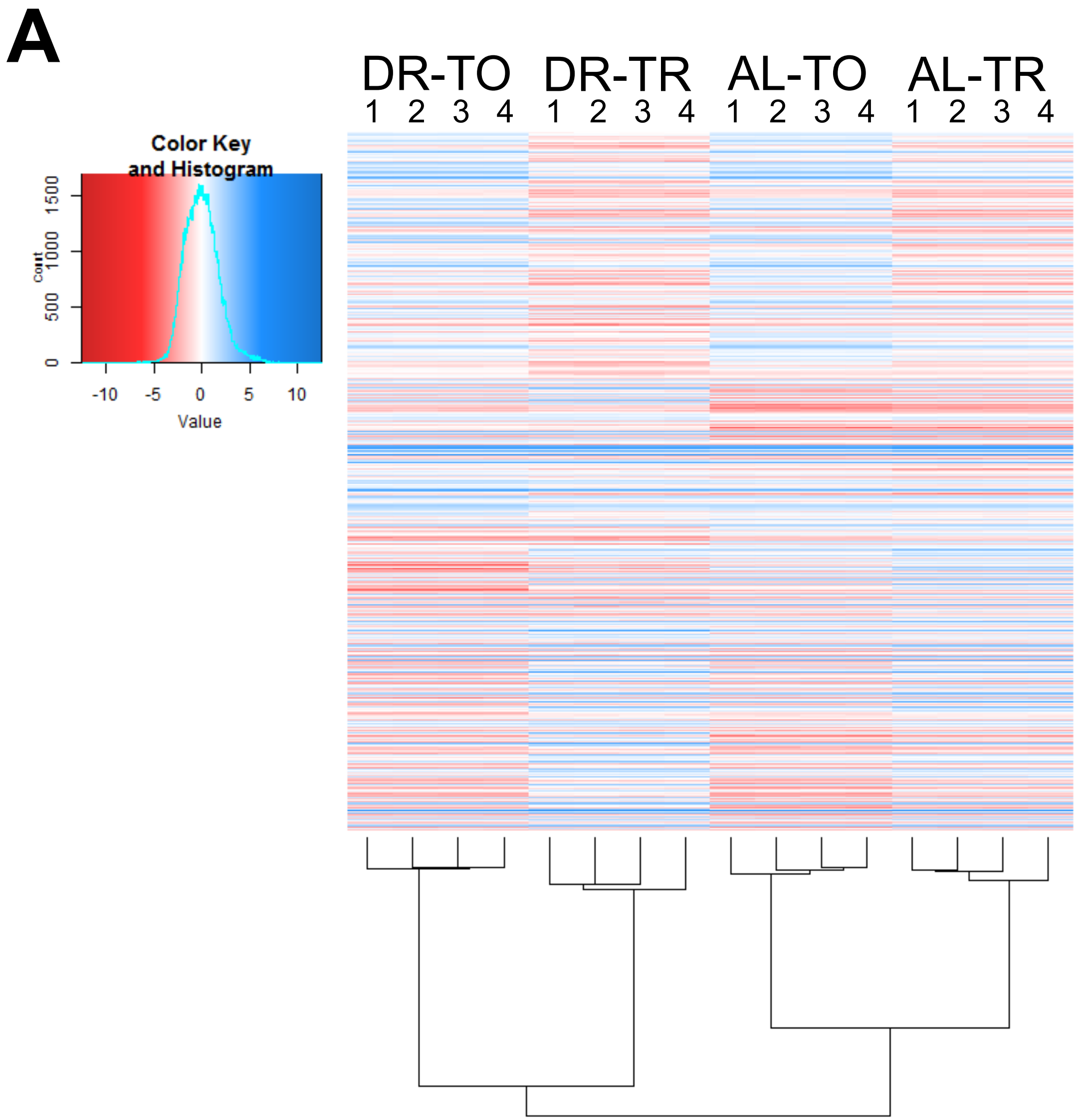


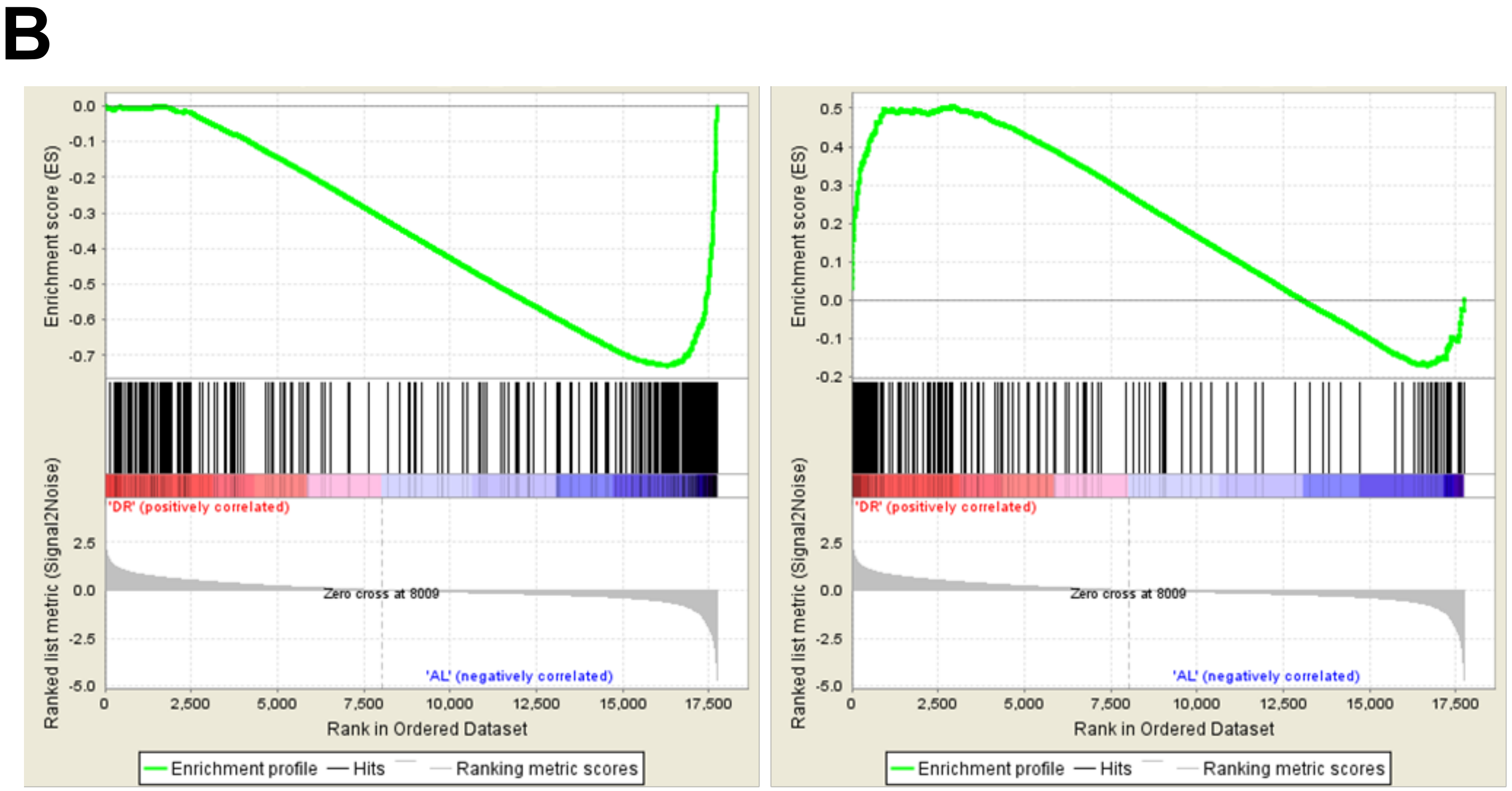


| | |
|---|---|
| Enrichment Score (ES) | -0.72 |
| Normalized Enrichment Score (NES) | -3.25 |
| FDR q-value | < 0.001 |

| | |
|---|---|
| Enrichment Score (ES) | 0.50 |
| Normalized Enrichment Score (NES) | 2.10 |
| FDR q-value | < 0.001 |

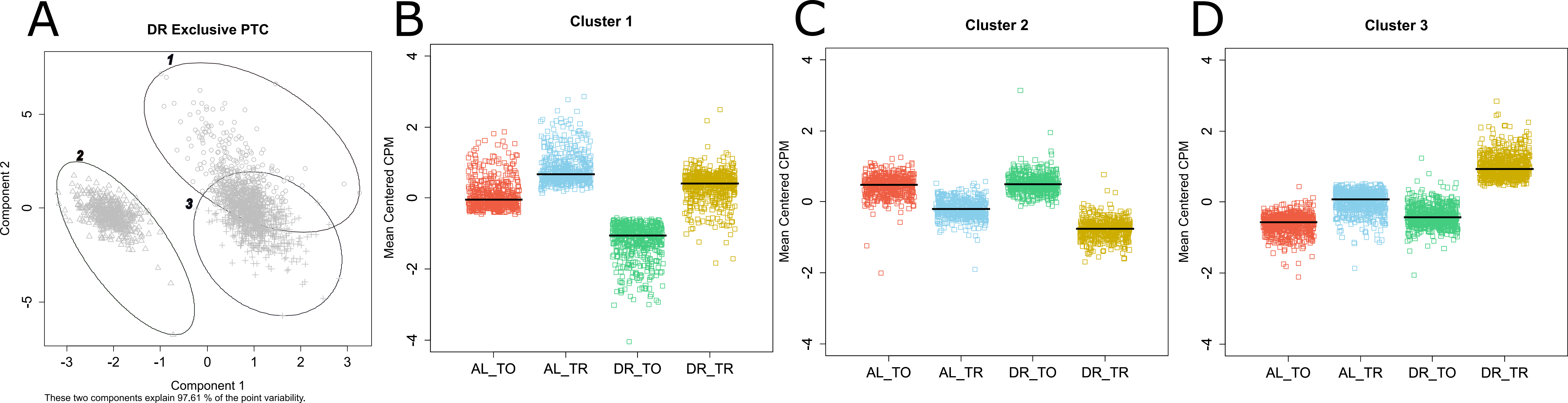
A
DR Exclusive PTC
1
2
3
Component 2
Component 1
These two components explain 97.61 % of the point variability.
B
Cluster 1
C
Cluster 2
D
Cluster 3
Mean Centered CPM
AL_TO
AL_TR
DR_TO
DR_TR

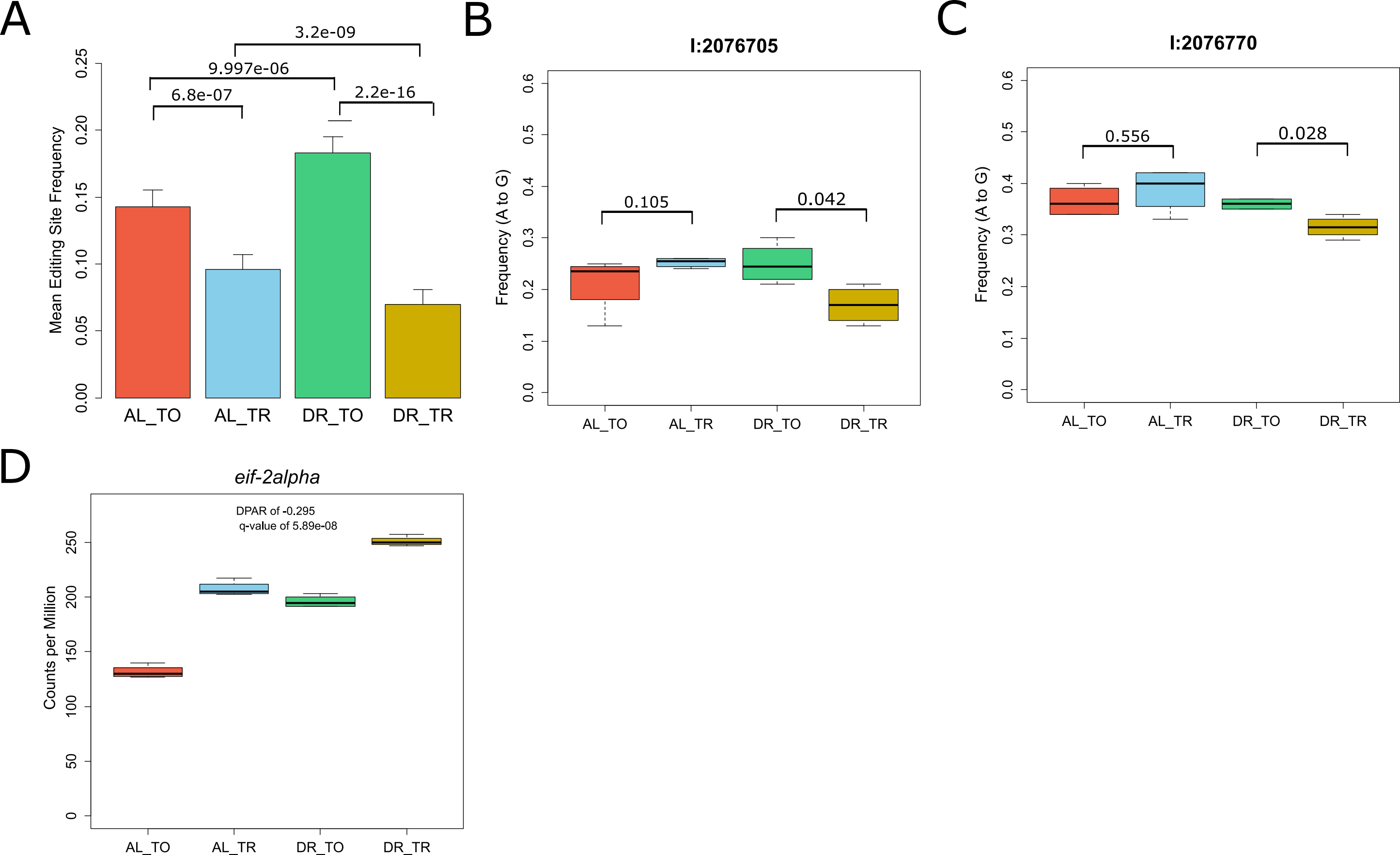
A
Mean Editing Site Frequency
0.00
0.05
0.10
0.15
0.20
0.25
3.2e-09
9.997e-06
6.8e-07
2.2e-16
AL_TO
AL_TR
DR_TO
DR_TR
B
I:2076705
Frequency (A to G)
0.0
0.1
0.2
0.3
0.4
0.5
0.6
0.105
0.042
AL_TO
AL_TR
DR_TO
DR_TR
C
I:2076770
Frequency (A to G)
0.0
0.1
0.2
0.3
0.4
0.5
0.6
0.556
0.028
AL_TO
AL_TR
DR_TO
DR_TR
D
eif-2alpha
Counts per Million
0
50
100
150
200
250
DPAR of -0.295
q-value of 5.89e-08
AL_TO
AL_TR
DR_TO
DR_TR

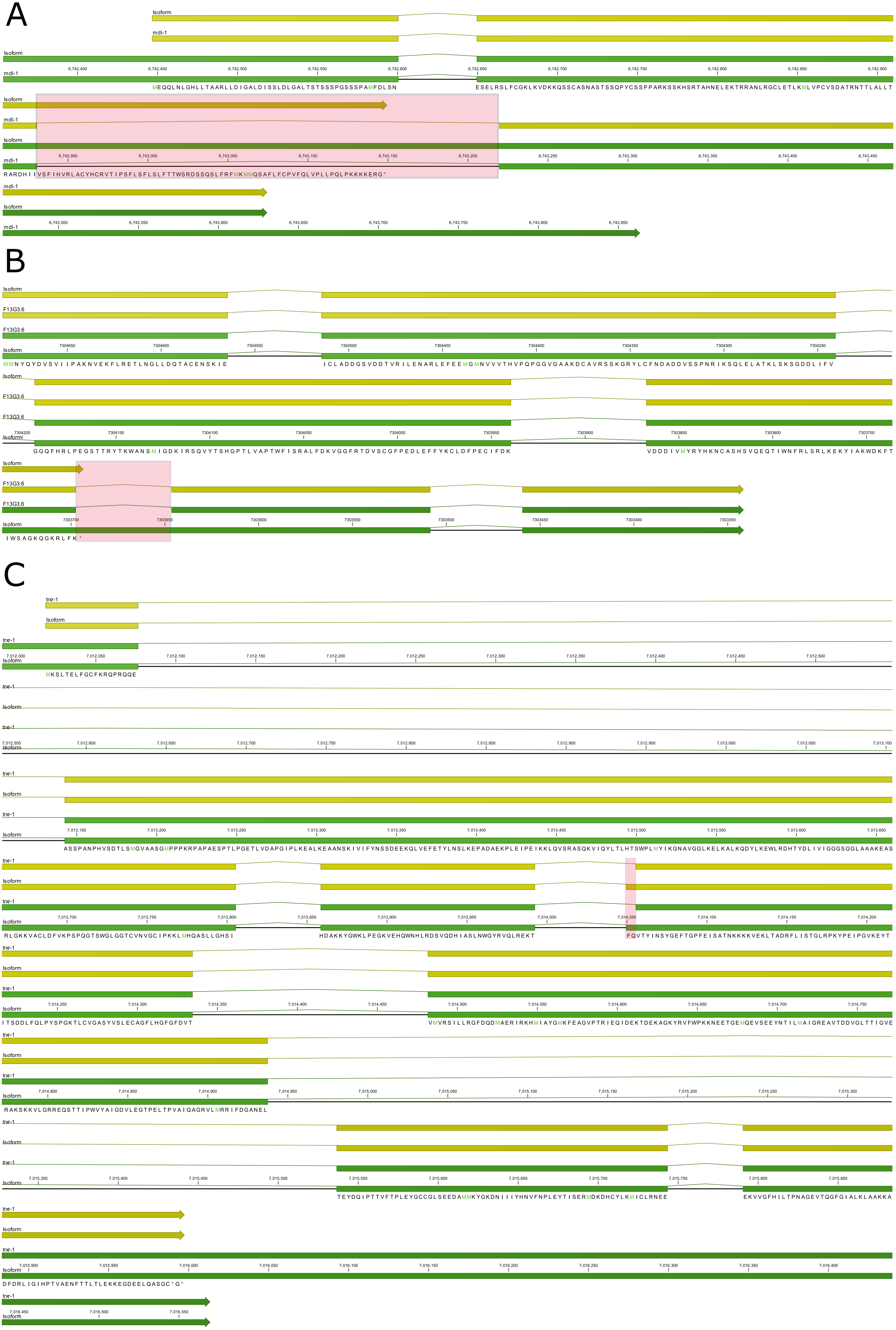
A
Isoform
mdl-1
B
Isoform
F13G3.6
C
trxr-1
Isoform
